# Modulation of type I interferon responses potently inhibits SARS-CoV-2 replication and inflammation in rhesus macaques

**DOI:** 10.1101/2022.10.21.512606

**Authors:** Timothy N. Hoang, Elise G. Viox, Amit A. Upadhyay, Zachary Strongin, Gregory K. Tharp, Maria Pino, Rayhane Nchioua, Maximilian Hirschenberger, Matthew Gagne, Kevin Nguyen, Justin L. Harper, Shir Marciano, Arun K. Boddapati, Kathryn L. Pellegrini, Jennifer Tisoncik-Go, Leanne S. Whitmore, Kirti A. Karunakaran, Melissa Roy, Shannon Kirejczyk, Elizabeth H. Curran, Chelsea Wallace, Jennifer S. Wood, Fawn Connor-Stroud, Sudhir P. Kasturi, Rebecca D. Levit, Michael Gale, Thomas H. Vanderford, Guido Silvestri, Kathleen Busman-Sahay, Jacob D. Estes, Monica Vaccari, Daniel C. Douek, Konstantin M.J. Sparrer, Frank Kirchhoff, R. Paul Johnson, Gideon Schreiber, Steven E. Bosinger, Mirko Paiardini

**Author notes:** Correspondence to (M.P; Lead Contact); (S.E.B.). These authors contributed equally (Timothy N. Hoang, Elise G. Viox, Amit A. Upadhyay). Deceased (Timothy N. Hoang).

## Abstract

Type-I interferons (IFN-I) are critical mediators of innate control of viral infections, but also drive recruitment of inflammatory cells to sites of infection, a key feature of severe COVID-19. Here, and for the first time, IFN-I signaling was modulated in rhesus macaques (RMs) prior to and during acute SARS-CoV-2 infection using a mutated IFNα2 (IFN-modulator; IFNmod), which has previously been shown to reduce the binding and signaling of endogenous IFN-I. In SARS-CoV-2-infected RMs, IFNmod reduced both antiviral and inflammatory ISGs. Notably, IFNmod treatment resulted in a potent reduction in (i) SARS-CoV-2 viral load in Bronchoalveolar lavage (BAL), upper airways, lung, and hilar lymph nodes; (ii) inflammatory cytokines, chemokines, and CD163+MRC1-inflammatory macrophages in BAL; and (iii) expression of Siglec-1, which enhances SARS-CoV-2 infection and predicts disease severity, on circulating monocytes. In the lung, IFNmod also reduced pathogenesis and attenuated pathways of inflammasome activation and stress response during acute SARS-CoV-2 infection. This study, using an intervention targeting both IFN-α and IFN-β pathways, shows that excessive inflammation driven by type 1 IFN critically contributes to SARS-CoV-2 pathogenesis in RMs, and demonstrates the potential of IFNmod to limit viral replication, SARS-CoV-2 induced inflammation, and COVID-19 severity.

## Main

Coronavirus disease 2019 (COVID-19) is caused by severe acute respiratory syndrome coronavirus 2 (SARS-CoV-2) and is an ongoing and rapidly developing pandemic^1^. While vaccines are highly effective, vaccine-induced immunity and neutralizing antibody titers wane over time. In addition, the emergence of SARS-CoV-2 variants resulting in breakthrough infections remains worrisome. Thus, it is imperative to fully characterize the viral pathogenesis of SARS-CoV-2 and early COVID-19 immune responses to identify correlates of protection and design therapeutics that can mitigate disease severity and viral replication.

Type-I interferons (IFN-I) are ubiquitously expressed cytokines that play a central role in innate antiviral immunity and cell-intrinsic immunity against viral pathogens^2–4^. The receptors for IFN-I, IFNAR1 and IFNAR2, are universally expressed and trigger the transcription of interferon stimulated genes (ISGs), that mediate a myriad of antiviral effector functions and promote recruitment of inflammatory cells^5^. Work early in the pandemic indicated that the IFN-I system has a protective effect from disease in SARS-CoV-2 infection, as individuals with severe COVID-19 were demonstrated to be more likely to have deficiencies to IFN-I responses, either by the presence of rare inborn errors (*TLR3*, *IRF7*, *TICAM1*, *TBK1*, or *IFNAR1*), neutralizing auto-antibodies against IFN-I, or the lack of production of IFN-I^6–15^. Additionally, Genome Wide Association Studies (GWAS) of critically ill COVID-19 patients have identified multiple critical COVID-19-associated variants in genes that are involved in IFN-I signaling including *IFNA10*, *IFNAR2*, *TYK2,* as well as type III interferon signaling *IL10RB*^16^. These findings contributed to the establishment of a model in which sustained IFN-I responses were deemed to be critical for protecting SARS-CoV-2-infected patients from progressing to severe COVID-19.

However, more recent work has suggested that the role of the IFN-I system in SARS-CoV-2 infection may be more complex than initially thought. In contrast to the aforementioned studies, Povysil et al. found no associations between rare loss-of-function variants in IFN-I and severe COVID-19^17^. Further, Lee et al. demonstrated that hyper-inflammatory signatures characterized by IFN-I response in conjunction with TNF/IL-1β were associated with severe COVID-19 in SARS-CoV-2 infected individuals^18^ and Blanco-Melo et al. showed that high IFN-I expression in the lung was linked to increased morbidity early in the COVID-19 pandemic^19^. Additionally, prolonged IFN signaling in murine models of respiratory infection have recently been shown to interfere with repair in lung epithelia, increasing disease severity and susceptibility to bacterial infections^20, 21^. Further work also has demonstrated the association of multiple ISGs with increased SARS-CoV-2 infection. Specifically, interferon-induced transmembrane proteins (IFITM 1-3) have been shown to be commandeered by SARS-CoV-2 to increase efficiency of viral infection^22^, the IFN-I-inducible receptor sialic acid-binding Ig-like lectin 1 (Siglec-1/CD169) can function as an attachment receptor and enhances ACE2-mediated infection and induction of trans-infection^23^, and prolonged cGAS-STING signaling has also been shown to induce aberrant IFN-I induced immunopathology by macrophages surrounding infected endothelial cells^24^. Lastly, IFN-I can also drive the expansion of pro-inflammatory monocytes (CD14^+^ CD16^+^), which have been shown to phagocytose FCγRs followed by abortive infection and induction of pyroptosis that induces systemic inflammation^25^.

Several ongoing and recently completed clinical trials administering IFNs have shown little to no positive effects of therapy during acute infection, despite treatment with IFN being highly efficient against SARS-CoV-2 *in vitro*^26, 27^. No beneficial effects were observed with either IFNβ-1a or combination IFNβ-1a and remdesivir treatment in in hospitalized COVID-19 patients in the WHO Solidarity Trial and Adaptive COVID-19 Treatment Trial (ACTT-3) respectively. In fact, combination IFNβ-1a and remdesivir actually resulted in worse outcomes and respiratory status in ACTT-3 patients who required high-flow oxygen at baseline when compared to treatment with remdesivir alone^28^. Pegylated IFN-α2b has shown some promise in moderate COVI-19 patients, with a phase 3 randomized open label study showing that patients that received pegylated IFN-α2b experienced a faster time to viral clearance^29^. Recently, there has been renewed interest in pursuing IFN-λ as a COVID-19 therapeutic following a recent initial release of the data on the Phase 3 TOGETHER clinical trial showing that early administration of pegylated IFN-λ resulted in a 50% risk reduction of COVID-19-related hospitalizations or lengthy emergency room visits in individuals at high risk of developing severe COVID-19^30^.

Given the discordant effects of IFN treatment observed in the aforementioned trials as well as the demonstrated ability of Type I and III IFNs to inhibit lung tissue repair in viral respiratory infections and render the host susceptible to opportunistic bacterial infections^20, 21^, it is critical that further research on therapies targeting IFN pathways is conducted to determine the optimal balance of stimulation of antiviral response while avoiding excessive inflammation and maintaining tissue repair pathways.

One compound previously used to effectively manipulate the IFN-I system is a mutated IFNα2 which binds with high affinity to IFNAR2, but markedly lower affinity to IFNAR1, with a net effect of reducing the binding and signaling of all forms of endogenous IFN-I^31–33^ (as such, previously referred to as IFN-I competitive antagonist, IFN-1ant). This mutated IFNα2, here named IFN-I modulator (IFNmod), has been shown to induce low-level stimulation of antiviral genes without induction of inflammatory genes when used *in vitro* in cancer cells^31, 32^ and, importantly, to limit the expression of both antiviral and pro-inflammatory ISGs when used *in vivo* in Simian Immunodeficiency Virus (SIV)-infected rhesus macaques (RMs)^33, 34^.

Non-human primate (NHP) models, specifically RMs, have been used extensively to study pathogenesis and evaluate potential vaccine and antiviral candidates for numerous viral diseases, including HIV and, more recently, SARS-CoV-2^35–38^. RMs infected with SARS-CoV-2 develop mild to moderate disease, mimic patterns of viral shedding, and, in similar fashion to humans, rarely progress to severe disease^35–38^. Multiple previous studies^35–38^ have shown that RMs generate a rapid and robust IFN-I response following SARS-CoV-2 infection, with numerous ISGs upregulated as early as 1-day post-infection. To better understand the role of IFN-I in SARS-CoV-2 infection, we characterized the effects of IFNmod on ISGs, SARS-CoV-2 viral replication, and pathology in an RM model.

## Results

### IFNmod administration decreases SARS-CoV-2 loads in airways of RMs

We first administered IFNmod to four uninfected RMs (7-10 years old, median = 9.5 years, **Supplementary Table 1**) at a dose of 1 mg/day, intramuscularly, for 4 consecutive days (**Supplementary Fig. 1a**) to determine its impact on ISGs in the absence of SARS-CoV-2 infection. Transcript levels of a panel of 17 ISGs previously observed to be induced by SARS-CoV-2 infection^35^ in RMs were quantified by RNAseq at pre- and post-treatment timepoints. ISGs were observed to be modestly upregulated following IFNmod administration in both BAL and PBMC of uninfected RMs (**Supplementary Fig. 1b-c**), whereas IL-6 signaling and inflammatory genes remained unchanged (**Supplementary Fig. 1d-e)**. Additionally, in concordance with the unchanged expression of IL-6 signaling and inflammatory genes, IFNmod treatment in uninfected animals did not impact cytokines and chemokines associated with inflammation and recruitment of inflammatory cells including IL-1β, IL-6, IL-8, IL-12p40, TNFβ, IFNγ, MIP1α, and MIP1β **(Supplementary Fig. 1f)**. Consistent with the data in uninfected RMs, IFNmod treatment alone in Calu-3 human lung cancer cells induced low-level expression of antiviral ISGs OAS1 and ISG15 but did not lead to detectable differences of the pro-inflammatory chemokine CXCL10 **(Supplementary Fig. 2a-c).** Importantly, in the presence of IFNα stimulation, IFNmod inhibited expression of both antiviral ISGs (ISG15, OAS1) and the pro-inflammatory CXCL10 **(Supplementary Fig. 2a-c)**. Taken together, these data indicate that IFNmod stimulates a weak antiviral IFN-I response in the absence of endogenous IFNα, but potently inhibits both antiviral and pro-inflammatory IFN-I pathways induced in response to addition of IFNα. In a follow up experiment, Calu-3 cells were treated with IFNmod and, 24-hours later, infected with SARS-CoV-2 NL-020-2020 (MOI 0.1) **(Supplementary Fig. 2d)**. qRT-PCR analysis of the supernatant at 48-hours post-infection demonstrated that IFNmod inhibited SARS-CoV-2 replication in a dose-dependent manner, with up to 80% reduction in viral RNA copies at the highest tested dose **(Supplementary Fig. 2e-f)**.

To determine how IFN-I pathways affect SARS-CoV-2 replication and pathogenesis *in vivo*, 18 adult RMs (6-20 years old, mean = 10 years; 10 males and 8 females; **Supplementary Table 1**) were age and sex matched between two experimental arms to receive 4 doses of IFNmod (intramuscularly, 1 mg/day; IFNmod-treated group) starting from one day prior to infection (d-1) and continued until 2 days post-infection (dpi), mimicking the 4-day dosing regimen of the uninfected RMs, or to remain untreated (untreated group) (**Fig. 1a**). On day 0, all 18 RMs were inoculated with a total of 1.1×10^6^ PFU SARS-CoV-2 (2019-nCoV/USA-WA1/2020), administered by intranasal (IN) and intratracheal (IT) routes. 3 IFNmod and 3 untreated RMs were euthanized at 2, 4, and 7 dpi each, respectively. The majority of animals displayed no overt clinical signs of disease following infection **(Supplementary Fig. 3a-b, Supplementary Tables 2 and 3)**. Overall, IFNmod was well tolerated without evidence of treatment induced clinical-pathology, nephrotoxicity, or hepatotoxicity when compared to untreated SARS-CoV-2 infected RMs.

**Fig. 1.**
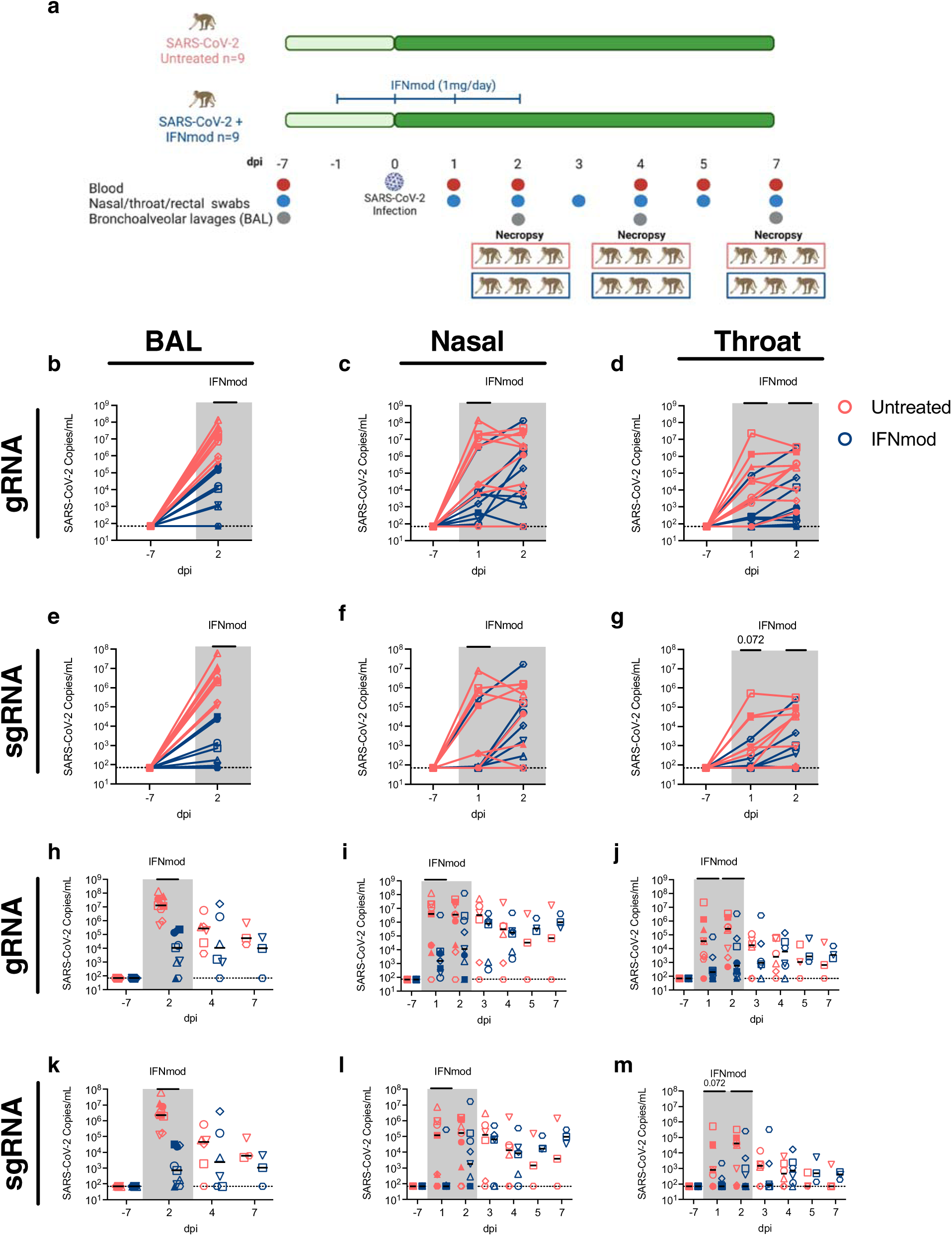
IFNmod administration results in significant reduction in viral loads of SARS-CoV-2 infected RMs. **(a)** Study Design; 18 RMs were infected intranasally and intratracheally with SARS-CoV-2. 1 day prior to infection (−1dpi), 9 RMs started a 4-dose regimen of IFNmod (1mg/day) that continued up until 2dpi while the other 9 RMs remained untreated. RMs were sacrificed at 2, 4, or 7 dpi. Levels of SARS-CoV-2 gRNA N **(b-d)** and sgRNA E **(e-g)** in BAL, nasopharyngeal swabs, and throat swabs during IFNmod treatment. Levels of SARS-CoV-2 gRNA N **(h-j)** and sgRNA E **(k-m)** in BAL, nasopharyngeal swabs, and throat swabs longitudinally throughout the entire duration of the study. Untreated animals are depicted in red and IFNmod-treated animals are depicted in blue. Black lines in h-m represent median viral loads for each treatment group at each timepoint. Gray-shaded boxes indicate that timepoint occurred during IFNmod treatment. Statistical analyses were performed using non-parametric Mann-Whitney tests. * p-value < 0.05, ** p-value < 0.01.

Viral RNA levels were measured using genomic (gRNA) and sub-genomic (sgRNA) qRT-PCR as previously described^35, 39, 40^. During the treatment phase of the study, up to 2 dpi, a drastic reduction in the levels of gRNA **(Fig. 1b-d)** and particularly sgRNA E **(Fig. 1e-g)** was observed between untreated and IFNmod-treated RMs in the BAL (median sgRNA E at 2 dpi: 2.31×10^6^ vs 7.18×10^2^, p < 0.0001 **Fig. 1e)**, nasal swabs (median sgRNA E at 1 dpi: 1.19×10^5^ vs 7.00×10^1^, p = 0.0123 **Fig. 1f**), and throat swabs (median sgRNA E at 1 dpi: 8.26×10^2^ vs 7.00×10^1^, p = 0.072; median sgRNA E at 2 dpi: 4.00×10^4^ vs 7.00×10^1^, p = 0.0496; **Fig. 1g)**. Of note, this corresponds to IFNmod treatment resulting in a >3000-fold reduction in median sgRNA E in the BAL at 2 dpi (**Fig. 1e**). Additionally, in the nasal swabs at 1 dpi, there was a >1500-fold reduction in median sgRNA E copies in IFNmod-treated RMs (**Fig. 1f**). The sgRNA E levels detected at 1 and 2 dpi in the throat showed similar differences, 12-fold and 570-fold less median sgRNA E detected in IFNmod-treated RMs compared to untreated RMs at these respective timepoints (**Fig. 1g**). Once treatment was stopped, viral loads remained stable in the treated group up until 7 dpi, both for the genomic **(Fig. 1h-j)** and the sub-genomic RNA (**Fig. 1k-m)**. As consistently shown in previous studies^35–38^, viral loads decreased in untreated, SARS-CoV-2-infected RMs shortly after the early peak, with swab and BAL viral loads no longer being statistically different between IFNmod-treated and untreated RMs starting from 4 dpi. BAL and nasal swab sgRNA E viral loads were reproduced by an independent laboratory (**Supplementary Fig. 4a**) and further confirmed by sgRNA targeting the N gene **(Supplementary Fig. 4b)**.

In summation, and consistent with the *in vitro* data in Calu-3 cells, *in vivo* treatment with IFNmod resulted in a highly reproducible and drastic (1-3 log_10_) decrease in viremia in the airways of SARS-CoV-2-infected RMs.

### IFNmod treatment reduces lung pathology and soluble markers of inflammation in SARS-CoV-2 infected RMs

To assess lung damage following SARS-CoV-2 infection, 3 RMs from each treatment group were euthanized at 2, 4 and 7 dpi. At necropsy, sections of the cranial (upper) and caudal (lower) lung lobes were taken for virologic and pathologic analyses. Notably, IFNmod-treated RMs necropsied during the treatment phase at 2 dpi had lower viral gRNA and sgRNA levels in their upper and lower lungs as well as hilar LNs than untreated RMs (**Fig. 2a-b)**. Although the small sample size at 2 dpi (Untreated= 3 RMs, IFNmod= 3 RMs) limited statistical power, this difference was substantial, with a 1500-fold and 500-fold reduction in mean gRNA and 250-fold and 300-fold reduction in mean sgRNA E observed in the upper and lower lung segments respectively of untreated vs. IFNmod-treated RMs **(Fig. 2a-b)**. Consistent with the ability of IFNmod to reduce BAL viral loads, treated RMs displayed decreased expression of nucleocapsid in the lung by immunohistochemistry **(Supplementary Fig. 4c-e)**. Furthermore, the nucleocapsid staining was more diffuse in the untreated RMs, whereas IFNmod-treated RMs had small foci of infected cells. Levels of Mx1 were also lower in IFNmod-treated RMs **(Supplementary Fig. 4f-h)** as compared to untreated RMs, confirming a reduction in the expression of proteins regulated by IFN-I.

**Fig. 2.**
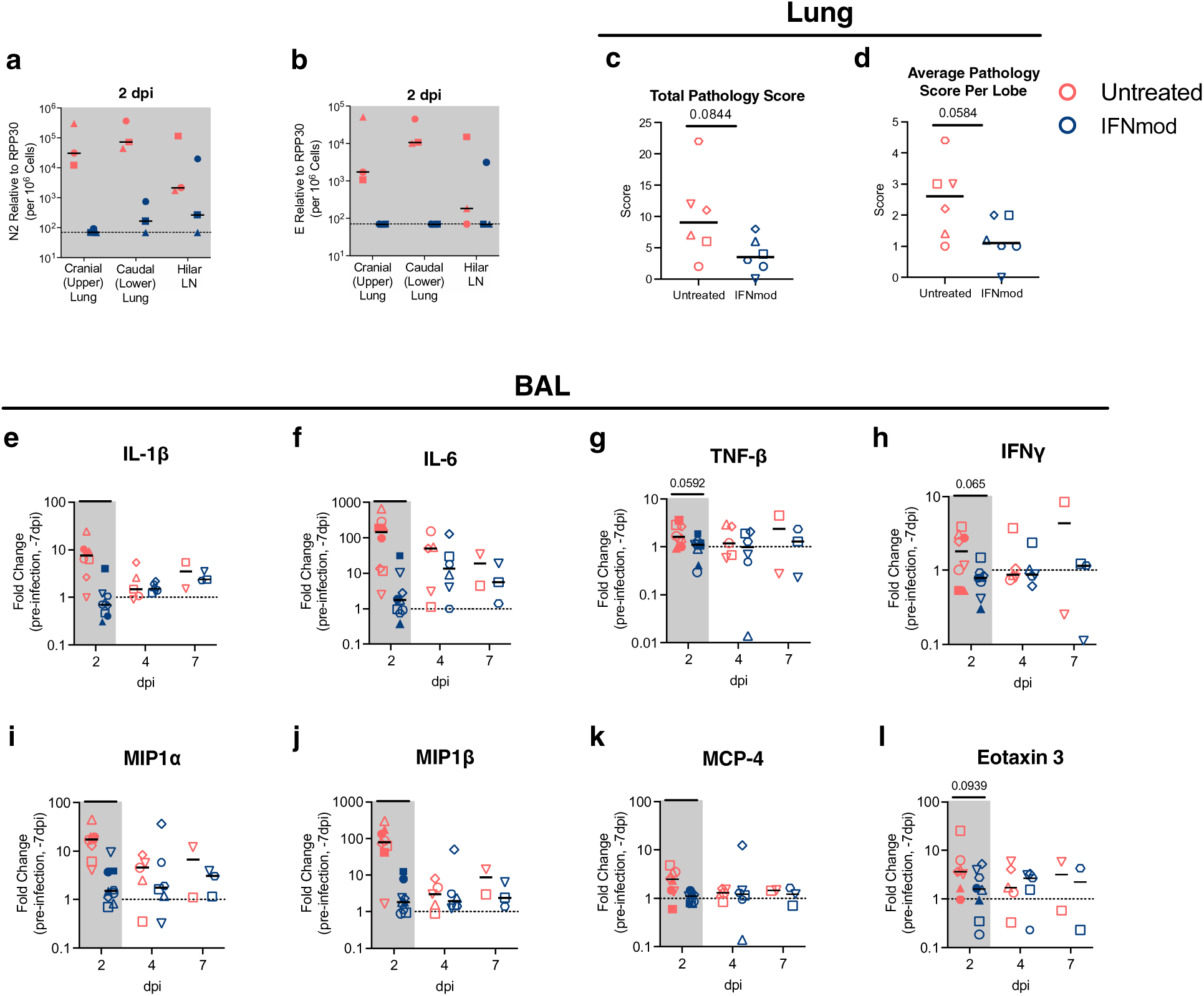
IFNmod administration resulted in lower levels of lung pathology and inflammatory cytokines and chemokines in SARS-CoV-2-infected RMs. Levels of SARS-CoV-2 gRNA **(a)** and **(b)** sgRNA in cranial (upper) lung, caudal (lower) lung, and hilar lymph nodes (LNs) of RMs necropsied at 2 dpi. **(c)** Total lung pathology scores and **(d)** average lung pathology scores per lobe of RMs necropsied at 4 and 7 dpi. **(e-l)** Fold change of cytokines and chemokines in BAL fluid relative to –7 dpi measured by mesoscale. Untreated animals are depicted in red and IFNmod treated animals are depicted in blue. Black lines represent the median viral load, pathology score, or fold change in animals from each respective treatment group. Gray-shaded boxes indicate that timepoint occurred during IFNmod treatment. Statistical analyses were performed using non-parametric Mann-Whitney tests. * p-value < 0.05, ** p-value < 0.01.

Pathological analysis of the lungs was performed as previously described^35^ by two pathologists, independently and blinded to the experimental arms using the following scoring criteria: type II pneumocyte hyperplasia, alveolar septal thickening, fibrosis, perivascular cuffing, and peribronchiolar hyperplasia. In both untreated and IFNmod-treated RMs necropsied at 2 dpi, no lung pathology was observed according to our scoring criteria. However, at 4 dpi and 7dpi, lung pathology was quantifiable, with untreated RMs having higher type 2 pneumocyte hyperplasia, alveolar septal thickening, and perivascular cuffing as compared to IFNmod-treated RMs **(Supplementary Fig. 4i**). The total lung pathology score (considering severity and number of effected lobes, p=0.0844) and average lung pathology score per lobe (measuring the average severity of abnormalities per lobe, independently of how many lobes had been affected, p=0.0584) were lower in the IFNmod-treated group (averages of 3.8 and 1.2, respectively) as compared to untreated RMs at 4 dpi and 7 dpi (averages of 10.0 and 2.5, respectively) (**Fig. 2c-d**).

A key feature of SARS-Cov-2-induced severe COVID-19 is the induction of multiple mediators of inflammation and chemotaxis of inflammatory cells to sites of infection^35^. Accordingly, multiple chemokines and cytokines were shown to be highly upregulated in the BAL of untreated RMs at 2 days post SARS-CoV-2 infection as measured by fold change (FC) to baseline (−7 dpi); remarkably, at 2 days post-infection the same cytokines and chemokines remained stable at basal levels in IFNmod-treated RMs (**Fig. 2e-l**). More specifically, we observed statistically significant differences in IL-1β (FC vs. pre-infection baseline at −7 dpi: 8.1 vs 0.81 respectively, p=0.001), IL-6 (FC: 196.9 vs 2.92, p=0.0016), MIP1α (FC: 16.93 vs 2.49, p=0.0003), MIP1β (FC: 105.2 vs 1.57, p=0.0025), and MCP4 (FC: 3.14 vs 1.02, p=0.0111) and trending differences in TNFβ (FC: 1.77 vs 0.99, p=0.0592), IFNγ (FC: 2.15 vs 0.86, p=0.065), and Eotaxin 3 (FC: 8.51 vs 2.32, p=0.0939) between untreated and IFNmod-treated RMs.

Altogether, IFNmod treatment potently reduced viremia and immune mediated pathogenesis in the lung, as well as the levels of multiple cytokines and chemokines in BAL that orchestrate the recruitment of inflammatory cells to sites of infection.

### IFNmod-treated RMs display decreased expansion of inflammatory monocytes and rapid downregulation of Siglec-1 expression

Several studies have reported an expansion of circulating inflammatory monocytes in blood of severe COVID-19 patients^41–43^. High-dimensional flow cytometry was performed on whole blood and BAL of RMs pre- and post-infection to quantify the immunological effects of IFNmod on the frequencies of classical (CD14^+^CD16^-^), non-classical (CD14^-^CD16^+^), and inflammatory (CD14^+^CD16^+^) monocytes as previously described (gating strategy depicted in **Supplementary Fig. 5)**.^35^. Representative staining of monocyte subsets in untreated and IFNmod-treated animals at pre- and post-infection timepoints is shown in **Fig. 3a**. The frequency of CD14^+^CD16^+^ monocytes in blood increased from 11% to 31% of total monocytes from –7 dpi to 2 dpi (FC of 2.7) in untreated RMs, but only from 14% to 18% (FC of 1.3) in IFNmod-treated RMs **(Fig. 3b-c).** This difference between treatment groups was maintained at 4 dpi, where CD14+CD16+ monocytes accounted for 19% of total blood monocytes in untreated animals (FC of 1.7 compared to –7dpi) while only accounting for 10% of total blood monocytes in IFNmod-treated RMs (FC of 0.7 compared to –7dpi) **(Fig. 3b-c).** At both 2 dpi and 4 dpi, these differences were specific for blood, with no difference observed within the BAL. Therefore, treatment with IFNmod potently limited the expansion of inflammatory monocytes, thus reducing the potential for systemic and lower airway inflammation.

**Fig. 3.**
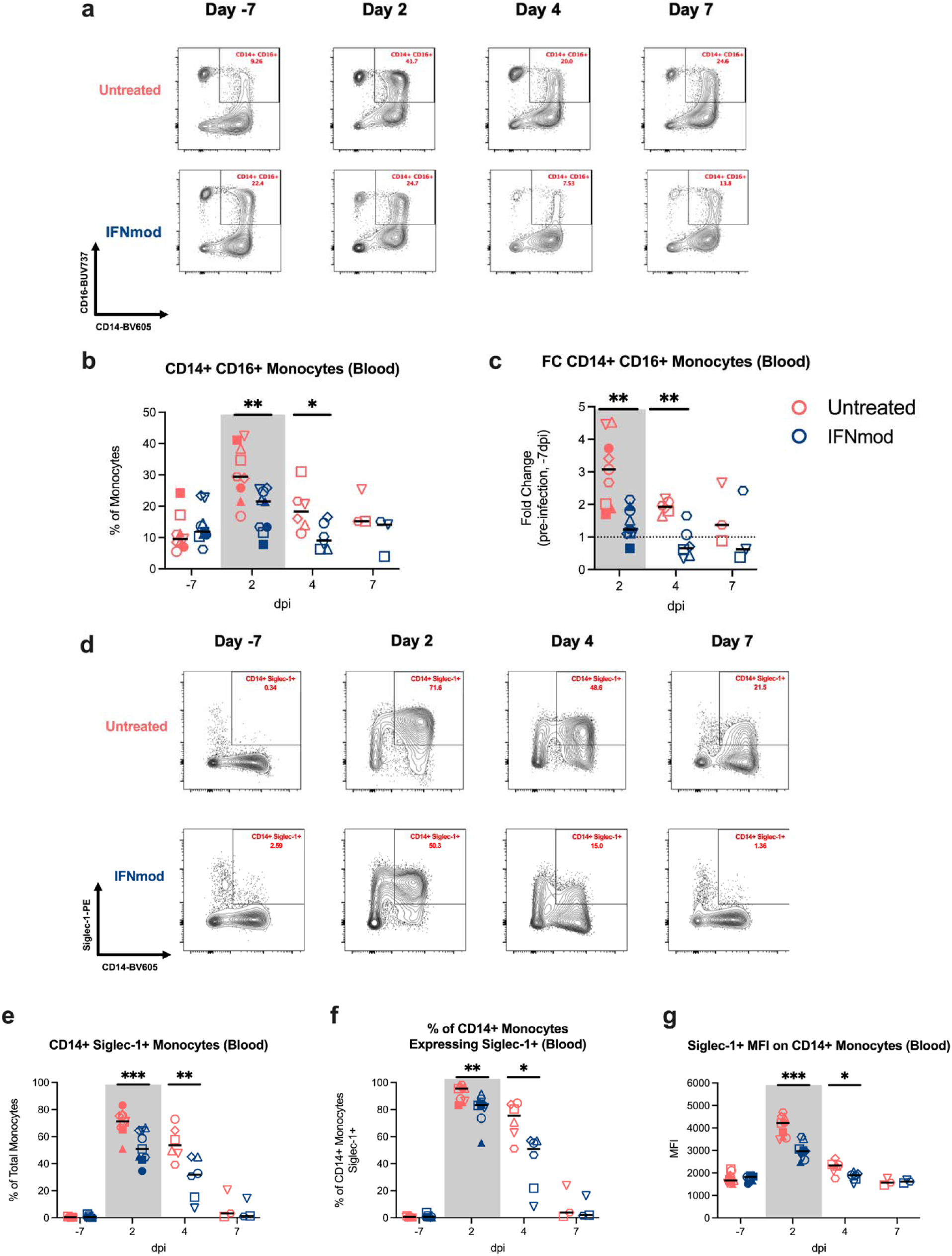
IFNmod-treated RMs had lower frequencies of CD14+CD16+ monocytes and expression of Siglec-1 compared to untreated RMs. **(a)** Representative staining of classical (CD14+CD16-), non-classical (CD14-CD16+), and inflammatory (CD14+CD16+) monocytes in peripheral blood throughout the course of infection. **(b-c)** Frequency and fold change relative to pre-infection baseline (−7dpi) of inflammatory (CD14+CD16+) monocytes in peripheral blood. **(d-e)** Representative staining and frequency of Siglec-1+ CD14+ monocytes in peripheral blood. **(f)** Frequency of CD14+ monocytes that were Siglec-1+ in peripheral blood. **(g)** MFI of Siglec-1 on CD14+ monocytes in peripheral blood. Untreated animals are depicted in red and IFNmod treated animals are depicted in blue. Black lines represent the median frequency or fold change in animals from each respective treatment group. Gray-shaded boxes indicate that timepoint occurred during IFNmod treatment. Statistical analyses were performed using non-parametric Mann-Whitney tests. * p-value < 0.05, ** p-value < 0.01

Siglec-1, an interferon responsive transmembrane protein present on antigen presenting cells, has been shown to function as an attachment receptor for SARS-CoV-2 through enhancement of ACE2-mediated infection and induction of trans-infection^23, 44^, and upregulation of Siglec-1 on circulating human monocytes has been identified as an early marker of SARS-CoV-2 infection and disease severity^45^. There was a strong and rapid upregulation of Siglec-1 on classical and inflammatory monocytes following SARS-CoV-2 infection in all untreated animals, with increases in the frequency of Siglec-1^+^CD14^+^ blood monocytes (as a percentage of total monocytes) from 0.4% to 70.6% (**Fig. 3d-e)**, frequency of CD14^+^ monocytes expressing Siglec-1 from 0.5% to 91.7% between –7 dpi and 2 dpi (**Fig. 3f),** and mean fluorescence intensity (MFI) of Siglec-1 on CD14^+^ monocytes from 1750 to 4050 (**Fig. 3g**). All 3 values remained significantly elevated at 4 dpi as compared to −7 dpi. Expression of Siglec-1 was significantly lower in the IFNmod-treated RMs when compared to untreated RMs both at 2 and 4 dpi. More specifically, IFNmod-treated RMs had a lower frequency of Siglec-1^+^CD14^+^ blood monocytes as a percentage of total monocytes (average at 2 dpi: 52.4% vs. 70.6%, p = 0.0008; average at 4 dpi: 29.3% vs. 55.2%, p = 0.0087; **Fig. 3d-e);** frequency of CD14^+^ monocytes expressing Siglec-1 (average at 2 dpi: 80.6% vs. 91.7%, p = 0.0078; average at 4 dpi: 41.0% vs. 72.2%, p = 0.0152; **Fig. 3f**); and MFI of Siglec-1 in CD14^+^ monocytes (average at 2 dpi: 3000 vs. 4050, p = 0.0003; average at 4 dpi: 1850 vs. 2250, p = 0.026; **Fig. 3g);**.

The reduced expression of Siglec-1, a well-established downstream molecule of interferon signaling, on blood monocytes in treated RMs is consistent with the observation that IFNmod attenuated antiviral and pro-inflammatory ISGs.

### IFNmod treatment attenuates inflammation and inflammasome activation in the lower airway following SARS-CoV-2 infection

To characterize the pathways that were impacted by IFNmod treatment *in vivo*, bulk RNA-Seq profiling of BAL, PBMCs, and whole blood was performed at multiple timepoints following SARS-CoV-2 infection. SARS-CoV-2 infection induced a robust upregulation of ISG expression in the BAL of untreated animals at 2 dpi. Notably, the transcript levels of ISGs in BAL were substantially attenuated in RMs treated with IFNmod relative to the levels in untreated animals at both 2 and 4 dpi (**Fig. 4a),** with 16/17 ISGs being significantly downregulated in IFNmod-treated RMs relative to untreated RMs at 2 dpi (**Fig. 4b and Supplementary Fig. 6a)**. By 7 dpi, ISG expression levels in the BAL of IFNmod-treated animals had returned to basal levels, whereas in untreated RMs, the ISGs in BAL remained significantly elevated (**Fig. 4a**). In PBMCs and whole blood, IFNmod-treated animals also had reduced expression of ISGs, inflammatory genes, and innate immune genes relative to untreated animals following infection **(Supplementary Fig. 6b-g)**.

**Fig. 4.**
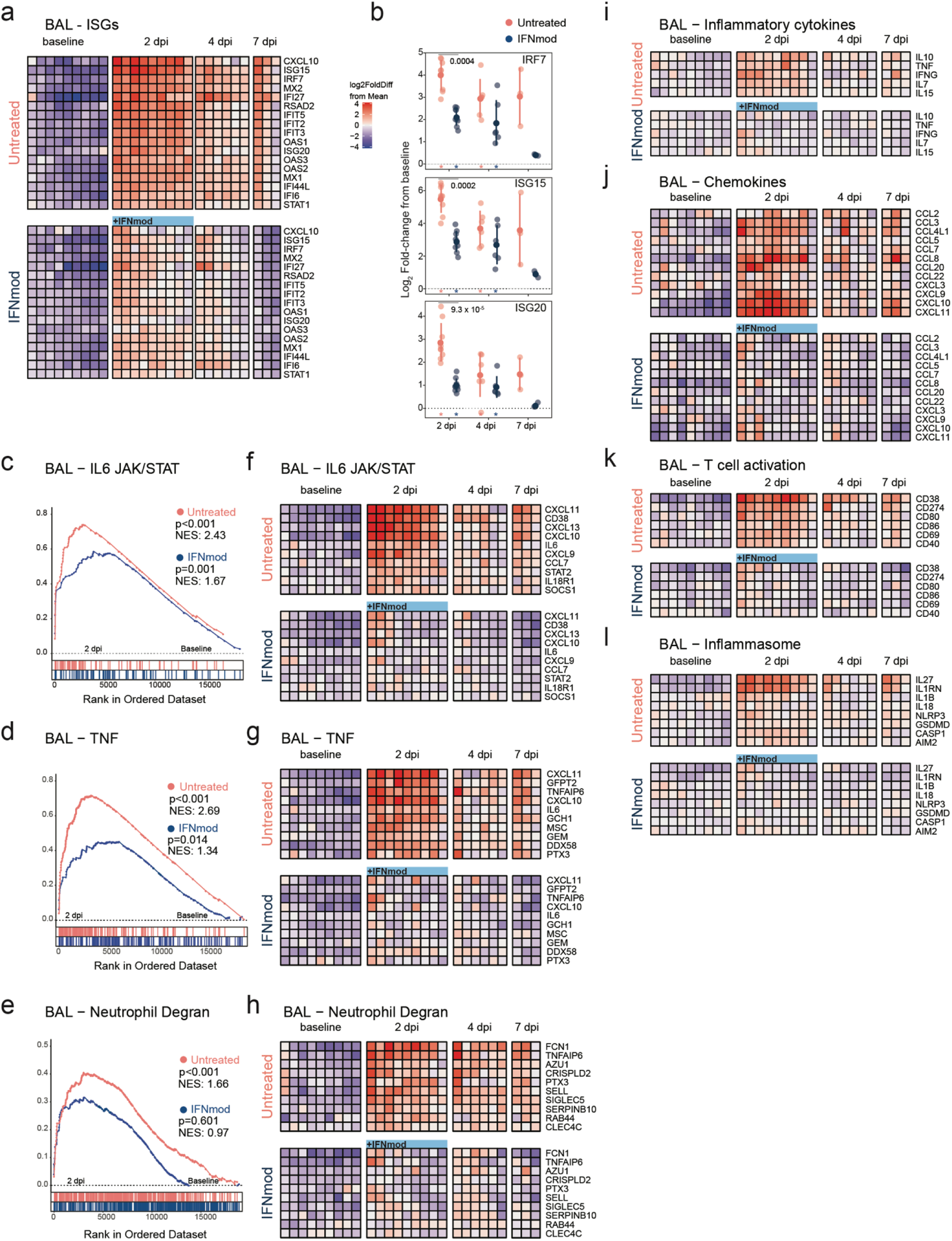
IFNmod treatment suppresses gene expression of ISGs, inflammation and neutrophil degranulation in the BAL of SARS-CoV-2 infected NHPs. Bulk RNA-Seq profiles of BAL cell suspensions obtained at −7 dpi (n = 9 for each arm), 2 dpi (n = 9 per arm), 4 dpi (n = 6 per arm), and 7 dpi (n = 3 per arm). **(a)** Heatmap of longitudinal gene expression in BAL prior to and following SARS-CoV-2 infection for the ISG gene panel. The color scale indicates log2 expression relative to the mean of all samples. Samples obtained while the animals were receiving IFNmod administration are depicted by a blue bar. **(b)** Distribution of log2 fold-changes of select ISGs relative to baseline. Filled dots represent the mean, and lighter dots are individual data points. Asterisks indicate statistical significance (p_adj_ < 0.05) of gene expression relative to baseline within treatment groups; black horizontal bars indicate p-values of direct contrasts of the gene expression between groups at time-points (i.e. IFNmod vs Untreated). **(c-e)** GSEA enrichment plots depicting pairwise comparison of gene expression of 2 dpi samples vs −7 dpi samples within treatment groups. The untreated group is depicted by red symbols, and data for the IFNmod treated group is shown in blue. The top-scoring (i.e. leading edge) genes are indicated by solid dots. The hash plot under GSEA curves indicate individual genes and their rank in the dataset. Left-leaning curves (i.e. positive enrichment scores) indicate enrichment, or higher expression, of pathways at 2 dpi, right-leaning curves (negative enrichment scores) indicate higher expression at −7 dpi. Sigmoidal curves indicate a lack of enrichment, i.e. equivalent expression between the groups being compared. The normalized enrichment scores and nominal p-values testing the significance of each comparison are indicated. Genesets were obtained from the MSIGDB (Hallmark and Canonical Pathways) database. **(f-h)** Heatmaps of longitudinal gene expression after SARS-CoV-2. Genes plotted are the top 10 genes in the leading edge of geneset enrichment analysis calculated in panels c-e for each pathway in the Untreated 2 dpi vs −7 dpi comparison. **(i-l)** Longitudinal gene expression for selected DEGs in immune signaling pathways. The expression scale is depicted in the top right of panel A.

To assess the impact of IFNmod treatment on inflammation in BAL, the post-infection enrichment of several gene sets in inflammatory pathways, previously associated with severe SARS-CoV-2 infection, was examined at 2 dpi. There was a noticeable loss in enrichment in *IL6 JAK/STAT* (NES 1.67 vs 2.43) **(Fig. 4c)** and *TNF* (NES 1.34 vs 2.69) **(Fig. 4d)** pathways in IFNmod-treated RMs as compared to untreated RMs. This was even more pronounced for genes associated with neutrophil degranulation **(Fig. 4e),** where IFNmod-treated RMs had no enrichment between 2 dpi and baseline (NES 0.971, p=0.601) and untreated RMs had a statistically significant enrichment (NES 1.66, <0.001). Examination of the leading-edge genes in these pathways (**Fig. 4f-h),** and differentially expressed genes (DEGS) identified by *DESeq2* analysis (**Fig. 4i-l**), displayed a stark divergence in the levels of several inflammatory mediators between the IFNmod and untreated groups. Notably, upregulation of IL6 (**Fig. 4f**), TNFA-inducing protein 6 (TNFAIP6), PTX3/TNFAIP5, TNF (**Fig. 4g, i**), IL10, IFNG, and IL7 (**Fig. 4i**) was evident in BAL samples in untreated animals but largely absent in IFNmod-treated animals (**Fig. 4f-g, i**). In untreated animals, genes encoding components of azurophilic granules (AZU1/azurocidin 1, RAB44) were elevated after infection; however, this expression was abrogated in the IFNmod-treated group (**Fig. 4h**). A similar pattern of expression was observed for several chemokines involved in migration and recruitment of monocytes and macrophages (CCL2, CCL3, CCL4L1, CCL5, CCL7, CCL8, CCL22), neutrophils (CXCL3), and activated T cells (CXCL9, CXCL10, and CXCL11) (**Fig. 4j)**. Several genes regulating T cell activation and co-stimulation (CD38, CD69, CD274/PDL1, CD80, CD86, CD40) were upregulated after SARS-CoV-2 infection in untreated animals but minimally changed in IFNmod-treated RMs (**Fig. 4k)**. Lastly, consistent with prior reports in human and mice ^46–48^, the expression of several S100 acute phase proteins (S100A8, S100A9, S100A12) were elevated in BAL samples from untreated animals but were absent in IFNmod-treated RMs. Collectively, these data demonstrate that IFNmod treatment induces a near complete loss of inflammatory activation and a significant reduction in pathways associated with recruitment and activation of myeloid cells and neutrophils in the lower airway.

Previously, the NLRP3 inflammasome pathway was shown to be the main driver of IFN-I induced pathology in SARS-CoV-2-infected hACE2 mice and inhibition of NLRP3 was observed to attenuate lung tissue pathology^49^. To investigate if NLRP3 signaling was induced by SARS-CoV-2 infection in RMs, we characterized the enrichment of genes associated with the two phases of NLRP3 signaling: i) priming, in which cytokines and PRRs can upregulate inflammasome components such as NLPR3, CASP1, pro-IL1b and ii) activation, which consists of IL1β and IL18 release and pyroptosis^50^. Canonical genes in the NLRP3 inflammasome pathway including IL1B, IL1RN/IL1RA, GSDMD, and NLRP3 were significantly upregulated (all p_adj_ < 0.05) at 2 dpi in BAL samples from untreated animals (**Fig. 4l**). Additionally, AIM2 was observed to be significantly upregulated in untreated RM BAL at 2 dpi, consistent with previous findings of AIM2 inflammasome activation in the monocytes of patients with COVID-19^25^. In IFNmod-treated RMs, however, there was no discernible upregulation from baseline of these inflammasome-associated factors (**Fig. 4l**). Thus, IFNmod treatment in RMs was able to potently attenuate inflammation and inflammasome activation systemically and in the lower airway during SARS-CoV-2 infection.

### IFNmod treatment inhibits the accumulation and activation of CD163^+^MRC1^-^ inflammatory macrophages in the lower airway during SARS-CoV-2 infection

To survey the impact of IFNmod treatment on the landscape of immune cells in the lower airway, droplet-based sc-RNA-Seq was performed on cells obtained from BAL pre- and post-SARS-CoV-2 infection in both IFNmod and untreated animals. After quality filtering, 62,081 cells were clustered using Seurat, followed by more refined annotation with a curated set of marker genes **(Supplementary Fig. 7a-c)**. Consistent with prior studies^35, 51^, the large majority of cells were of myeloid lineage (macrophages and monocytes), with a minority of several other immune phenotypes and epithelial cells observed (**Fig. 5a, Supplementary Fig. 7d**). The expression of the 17 ISG panel utilized above was examined in these subtypes (except for neutrophils) and showed that ISGs peaked at 2 dpi in untreated animals in all subsets **(Supplementary Fig. 7e-g)**. Remarkably, IFNmod treatment diminished expression of those ISGs in all of the cell subsets identified in the BAL **(Supplementary Fig. 7e-g)**. Thus, IFNmod is highly effective at reducing the IFN-I response in all cells identified in the lower airway.

**Fig. 5.**
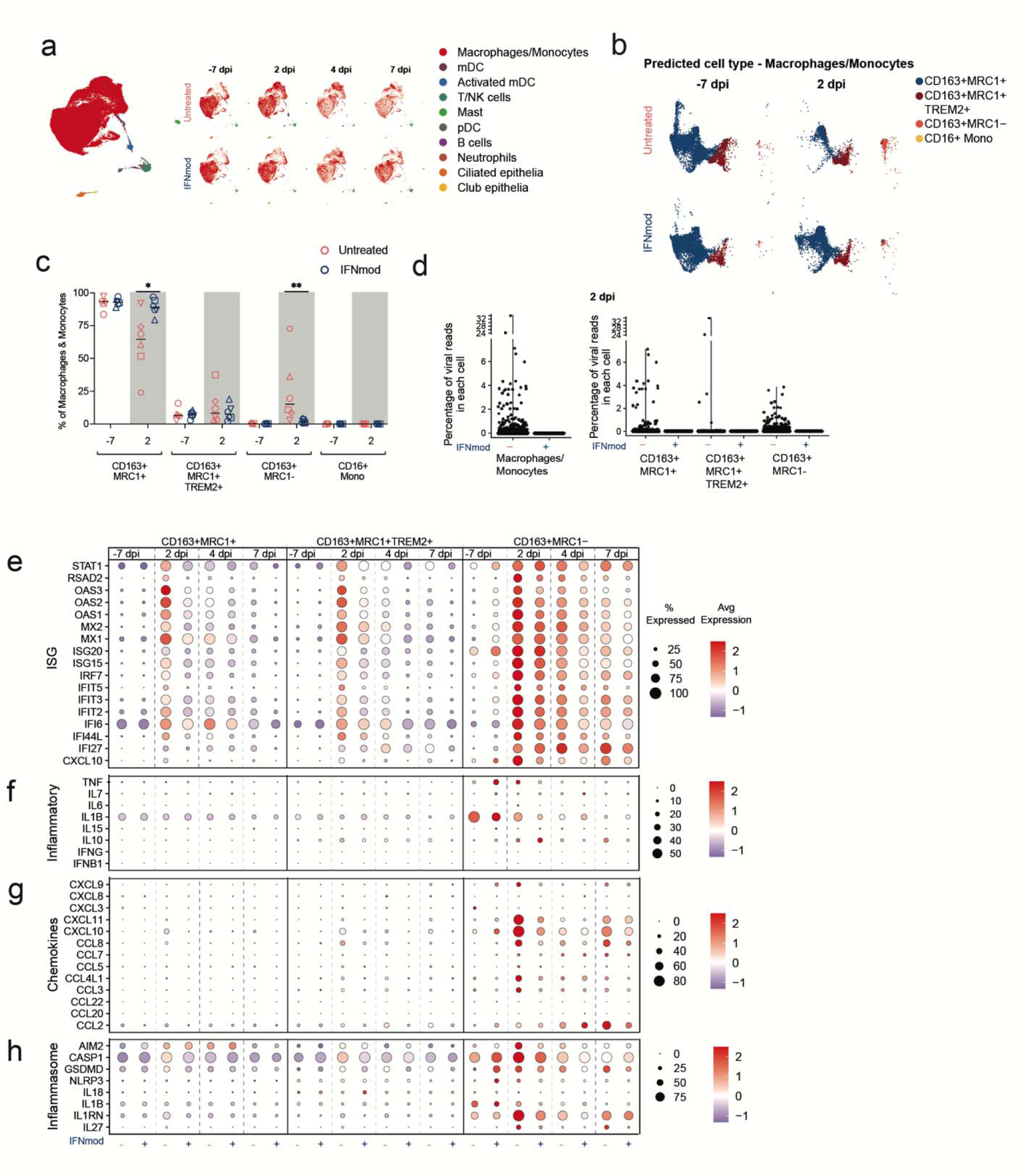
Effect of IFNmod treatment on gene expression of BAL single-cells using 10X. **(a)** UMAP of BAL samples (62,081 cells) integrated using reciprocal PCA showing cell type annotations. UMAP split by treatment and time points are also shown. **(b)** Mapping of macrophage/monocyte cells in the BAL of SARS-CoV-2-infected untreated and IFNmod treated RMs to different lung macrophage/monocyte subsets from healthy rhesus macaque ^91^. **(c)** Percentage of different macrophage/monocyte subsets out of all the macrophage/monocytes in BAL at −7 dpi and 2 dpi from untreated and IFNmod treated RMs. The black bars represent the median. Statistical analyses between treatment groups were performed using non-parametric Mann-Whitney tests. * p-value < 0.05, ** p-value < 0.01. **(d)** Violin Plots showing the percentage of viral reads in each cell for total macrophages/monocytes and individual macrophages/monocytes subsets at 2 dpi determined using the PercentageFeatureSet in Seurat. **(e-h)** Dot Plots showing expression of selected ISGs **(e),** inflammatory genes **(f),** chemokines **(g)** and inflammasome genes **(h).** The size of the dot indicates the percentage of cells that express a given a gene and the color indicates the level of expression. The numbers of CD16+ monocytes were very low and have been thus omitted.

In recent work, we used sc-RNA-Seq to identify two myeloid population of the BAL of RMs, CD163^+^MRC1^+^TREM2^+^ and CD163^+^MRC1^-^, that infiltrated into the lower airway and produced inflammatory cytokines during acute SARS-CoV2 infection. We also showed that blocking the recruitment of these subsets with a JAK/STAT inhibitor abrogated inflammatory signaling ^35, 52^. Here, using a reference comprising of the macrophage/monocyte subsets from lungs of three healthy RMs as described previously^52, 53^, we divided the clusters of Macrophage/Monocyte cells in the lower airways of RMs into four subsets: CD163^+^MRC1^+^, CD163^+^MRC1^+^TREM2^+^, CD163^+^MRC1^-^, and CD16^+^ monocytes (**Fig. 5b, Supplementary Fig. 8a-b**). Of note, the CD16+ monocyte subset was omitted from further functional analyses due to its very low frequency. The percentage of CD163^+^MRC1^+^ macrophages in the BAL of untreated RMs decreased following SARS-CoV-2 infection (median percentage of 93% at −7dpi to 65% at 2 dpi, p = 0.0625) due to the infiltration of CD163^+^MRC1^-^ and CD163^+^MRC1^+^TREM2^+^ cells at 2 dpi (**Fig. 5c, Supplementary Fig. 8c**). Conversely, the percentage of CD163^+^MRC1^+^ macrophages in the BAL of IFNmod-treated RMs remained relatively stable (median percentage of 93% at - 7dpi to 89% at 2 dpi), with CD163^+^MRC1^+^ macrophages at 2 dpi being significantly higher in IFNmod-treated vs. untreated RMs (p=0.0260). Additionally, the CD163^+^MRC1^-^ population in BAL was shown to have expanded significantly more from pre-infection baseline to 2 dpi in untreated animals (median percentages of 0.2% to 15%) as compared to IFNmod-treated animals (median percentages of 0.2% to 1.5%) (p = 0.0087) (**Fig. 5c, Supplementary Fig. 8c).** At 2 dpi, viral reads were detected (percentage of viral reads > 0) in macrophages/monocytes in the BAL of untreated RM at a rate of 9.8% (1030/9451 cells, median of individual animals 5.8%) compared to 0.01% (1/7544) in the BAL of IFNmod-treated RMs (**Fig. 5d, Supplementary Fig. 8d**). Among the macrophage subsets in the untreated group, the CD163+MRC1-subset was primarily associated with viral reads with a median of 11% of cells with detectable viral reads compared to 2.2% of CD163+MRC1+ and 2.4% of CD163+MRC1+TREM2+ (**Fig. 5d, Supplementary Fig. 8e**). The number of cells with detectable viral reads gradually decreased at 4dpi and was almost absent by 7 dpi **(Supplementary Fig. 8e)**.

Peak ISG levels in CD163^+^MRC1^+^, CD163^+^MRC1^+^TREM2^+^, and CD163^+^MRC1^-^ macrophage subsets in BAL were observed at 2 dpi, with the highest and most sustained ISG expression occurring in the CD163^+^MRC1^-^ population (**Fig. 5e**). In all three macrophage subsets in the BAL, IFNmod treatment effectively suppressed ISG expression (**Fig. 5e**). The CD163^+^MRC1^-^ were the predominant subset producing inflammatory cytokines and chemokines at 2 dpi and other post-infection timepoints in both experimental groups, with IFNmod reducing the transcript levels of both (**Fig. 5f-g**).

Since a series of studies in humans and a humanized mouse model linked lower airway inflammation during SARS-CoV-2 infection to inflammasome activation specifically within infiltrating monocytes and resident macrophages^25, 49^, we went on to characterize the expression of canonical inflammasome genes within each myeloid subset in the lower airway. We observed that expression of AIM2, CASP1, GSDMD, IL1B, IL1RN and IL27 increased from −7 dpi to 2 dpi in the CD163+MRC1-subset and was maintained until 7 dpi (**Fig. 5h**). Induction of expression for several inflammasome-associated genes was also observed at 2 dpi in CD163^+^MRC1^+^ and CD163^+^MRC1^+^TREM2^+^ macrophages, albeit at lower cellular percentages overall compared to the CD163^+^MRC1^-^ population **(Fig. 5h)**. Across all three myeloid subsets, IFNmod consistently dampened the expression of inflammasome mediators, with the effect being most apparent at 2 dpi **(Fig. 5h)**.

Collectively, these data indicate a profound effect of IFNmod treatment in inhibiting the accumulation of CD163^+^MRC1^-^ macrophages, the most prevalent subset undergoing inflammasome activation and contributing to inflammation, in the lower airway during SARS-CoV-2 infection.

### IFNmod treatment dampens inflammasome activation, bystander stress response, and cell death pathways in lung during acute SARS-CoV-2 infection

To further characterize the effect of IFNmod within the lower airway, sc-RNA-Seq was conducted on cell suspensions prepared from caudal (lower) lung lobe sections obtained at necropsy (2 dpi = 2 untreated, 2 IFNmod-treated; 7 dpi= 1 untreated, 1 IFNmod-treated). Based on the expression of canonical markers, these cells were classified into four major groups – Epithelial, Lymphoid, Myeloid and “Other”-stromal and endothelial (**Fig. 6a**), which were then each clustered separately. The annotations were further fine-tuned based on the expression of marker genes^54^ **(Supplementary Fig. 9a-d**), yielding 42,699 cells with balanced representation of varying phenotypes across the lung (**Fig. 6a and Supplementary Fig. 9a-d**). Interestingly, the Myeloid cluster in the lung contained an additional subset beyond the four phenotypes described in the BAL that was closely related to the CD163^+^MRC1^+^ cells and was defined by high expression of SIGLEC1 (**Fig. 6a and Supplementary Fig. 9c**).

**Fig. 6.**
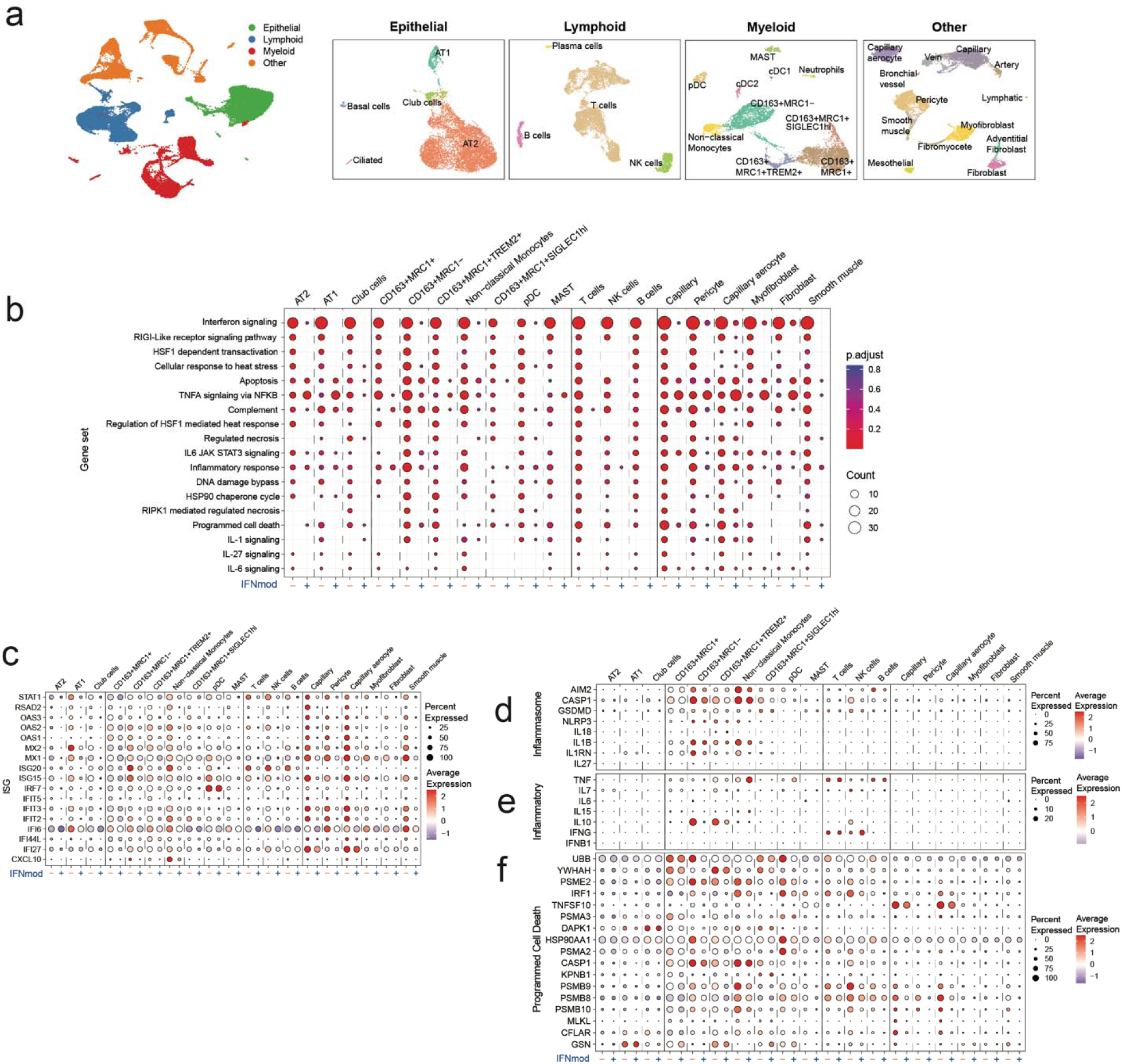
Effect of IFNmod treatment on lung cells. **(a)** UMAP based on reciprocal PCA of lung single cells collected at 2 dpi (n = 2 Untreated, 2 IFNmod) and 7 dpi (n = 1 Untreated, 1 IFNmod). The cells were classified into four broad categories – epithelial, lymphoid, myeloid and others (stromal and endothelial). The cells from each category were subset and clustered separately. UMAPs for each category with cell type annotations are also shown. **(b)** Selected gene sets that were found to be enriched (p-adjusted value < 0.05) in lung cells from untreated RMs at 2 dpi based on over-representation analysis using Hallmark, Reactome, KEGG, and BioCarta gene sets from msigdb. The size of the dots represents the number of genes that were enriched in the gene set and the color indicates the p-adjusted value. The gene set id in order are: M983, M15913, M27255, M27253, M5902, M5890, M5921, M27250, M41804, M5897, M5932, M27698, M27251, M29666, M27436, M27895, M27897, M1014. **(C-F)** Dot plots showing gene expression in lung cells present at higher frequencies from untreated and IFNmod treated macaques at 2 dpi **(c)** ISG, **(d)** genes related to inflammasome, **(e)** inflammation, and **(f)** programmed cell death. The size of the dot represents the percent of cells expressing a given gene and the color indicates the average expression.

Analyses at the pathway level at 2 dpi demonstrated a profound impact of IFNmod treatment in abrogating IFN-I signaling, as observed by an absence of enrichment of several IFN-related gene sets across all cell subsets in the lungs of IFNmod-treated RMs, despite being robustly overrepresented in untreated animals (**Fig. 6b and Supplementary Fig. 10a**). At the gene level, analyses of individual cell subsets, including AT1 and AT2 cells (Epithelial cluster), CD163^+^MRC1^-^ cells (Myeloid cluster), T cells, NK cells, and B cells (in the Lymphoid cluster) showed between 12 to 17 ISGs out of the panel of 17 ISGs being statistically significantly lower (p_adj_ < 0.05) in IFNmod-treated RMs as compared to untreated RMs at 2dpi (**Fig. 6c and Supplementary Fig. 10b**). At 7 dpi, five days after cessation of IFNmod treatment, expression of ISGs had largely normalized and was equivalent between the untreated and IFNmod groups (**Supplementary Fig. 10c**). Taken together, these data demonstrate that IFNmod treatment was able to effectively suppress the IFN-I system across a broad distribution of pulmonary cellular subsets during acute SARS-CoV-2 infection, including epithelium involved in gas exchange, immune cells in the interstitium, and cells in the lung vasculature.

Given the attenuated expression of genes related to inflammasome signaling in the BAL of IFNmod-treated RMs discussed previously, we investigated if a similar dampening effect occurred in the lungs of treated animals. In untreated animals, we observed a significant enrichment of genes within the IL1 signaling cascade at 2 dpi relative to pre-infection baseline across several myeloid, lymphocyte, and endothelial subsets. Conversely, in IFNmod-treated RMs, there was no enrichment of the IL1 signaling cascade geneset at 2dpi (**Fig. 6b)**. Additionally, there was a significant enrichment of genesets representing necrotic cell-death pathways observed in untreated RMs, but not IFNmod-treated RMs **(Fig. 6b).** Since infection-induced pyroptosis is a process of inflammation-related necrosis that has been shown to largely occur within myeloid cells^55^, we next delineated the contribution of individual subsets to the inflammatory state in the lower airway by visualizing DEGs across subsets (**Fig. 6d-f and Supplementary Fig. 10d-h**). Notably, expression of components of the inflammasome **(Fig. 6d and Supplementary Fig. 10e)** and classical inflammatory mediators **(Fig. 6e and Supplementary Fig. 10f)** were restricted to the myeloid subsets, suggesting that inflammasome priming and activation in the lung during SARS-CoV-2 infection is largely constrained to monocyte/macrophage populations. Consistent with this model, expression of GSDMD, a central regulator of NLRP3-regulated pyroptosis, was highest in monocyte/macrophage populations (**Fig. 6d**). Importantly, levels of these inflammasome components were all potently reduced by IFNmod treatment (**Fig. 6d**).

In untreated RMs, widespread induction of genes from the programmed cell death **(Fig. 6f and Supplementary Fig. 10g)** and response to heat stress (**Supplementary Fig. 10d, h**) pathways were observed across the landscape of lung cellular subsets, particularly in the Myeloid and Lymphoid classes of phenotypes. In contrast, within the IFNmod-treated group, expression of genes related to programmed cell death and response to heat stress was strikingly attenuated; this effect was particularly apparent in capillary and capillary aerocytes (**Fig. 6f, Supplementary Fig. 10d, g-h**).

These data are consistent with a model in which, during acute SARS-CoV-2 infection, activated monocytes and macrophages cells undergo inflammasome-mediated pyroptosis, which provides secondary stress signals to the non-immune cell classes of the lung. Importantly, IFNmod treatment was also able to abolish induction of these stress and cell death related pathways in these pulmonary subsets.

## Discussion

The ways in which IFN-I regulate SARS-CoV-2 replication and disease progression are incompletely understood. Here, we employed an integrated systems approach in a nonhuman primate model to dissect the opposing roles of antiviral and pro-inflammatory IFN-I responses in SARS-CoV-2 using a mutated IFNα2 (IFN-I modulator; IFNmod) that was previously demonstrated to have a high affinity to IFNAR2 and markedly lower affinity to IFNAR1, result in receptor occupancy, and block binding and signaling of all forms of endogenous IFN-I produced in response to viral infection^31–33^. While SARS-CoV-2-infection in untreated RMs was shown to induce a strong IFN-I response with concomitant inflammation and lung damage, IFNmod remarkably and potently reduced viral loads in the upper and lower airways, inflammation and lung pathology. Therefore, our study shows for the first time in NHP and with an *in vivo* intervention targeting IFN-I pathways that an aberrant IFN-I response critically contributes to SARS-CoV-2 pathogenesis, and demonstrates a beneficial role of IFNmod in limiting both viral replication and SARS-CoV-2 induced inflammation.

The administration of IFNmod prior to and during the first two days of SARS-CoV-2 infection was found to result in a highly significant reduction in viral loads in the BAL (>3 log reduction), nasal, and throat swabs. Additionally, IFNmod reduced viral loads in the upper and lower lungs as well as hilar LNs of RMs during the treatment period and limited lung pathology as well as the production of numerous pro-inflammatory cytokines and chemokines within the lower airways.

We show that IFNmod administration prior to SARS-CoV-2 infection in RMs results in a subtle upregulation of antiviral ISGs, while potently dampening endogenous IFN-I signaling and limiting the immunopathology that has been linked to IFN-I following SARS-CoV-2 infection. These data support a working model in which a sustained and systemic IFN-I response, particularly when extended to pro-inflammatory genes, exacerbate SARS-CoV-2 pathology and suggests that targeting downstream modulators of IFN-I responses may prove to be a viable therapeutic option. Of note, while IFNmod reduced viral loads and pathology in our SARS-CoV-2-infected RMs, it resulted in significantly higher plasma viral loads, increased CD4+ T cell depletion, and a more rapid progression to AIDS in SIV-infected RMs when administered during the first 4 weeks of infection. These results highlight that modulating IFN-I during infection with inherently different viruses (SARS-CoV-2, which establishes an acute infection that is naturally controlled in a few weeks in RMs and SIV, which establishes a persistent, chronic infection) can have vastly different effects on pathogenesis.

Our data also show a central role of IFN-I in regulating the axis of infiltrating monocytes and subsequent inflammation within the airway and lung interstitium during SARS-CoV-2 infection. Numerous studies have observed hyperactivation of monocytes and macrophages in COVID-19 (reviewed in^56^). Within the lower airway, an early study identified elevated expression of IL1B, IL6, TNF within alveolar macrophages in patients with severe COVID-19^51^. Subsequent work has progressed to dissect the precise phenotypes contributing to SARS-CoV-2 driven inflammation within the monocyte/macrophage axis in mice^57^, NHPs^37, 52^, and more recently, in humans^58^. In these studies, a consistent model has emerged in which resident tissue alveolar macrophages are displaced by inflammatory, infiltrating monocytes and interstitial macrophages. Notably, we observed IFNmod treatment to profoundly impact monocyte/macrophage populations *in vivo*, decreasing the activation state of the CD163^+^MRC1^-^ population and its recruitment to the BAL as well as reducing the expansion of CD14^+^CD16^+^ pro-inflammatory monocytes in the periphery relative to untreated RMs. We also identified the CD163^+^MRC1^-^ interstitial macrophage-like subset as the predominant population expressing inflammatory cytokines and chemokines and exhibiting inflammasome activation within the alveolar space during SARS-CoV-2 infection. Moreover, and supporting recent studies in humanized mice^49^, our work demonstrates that targeted modulation of the IFN-I system during early SARS-CoV-2 can effectively eliminate the level of cell-associated virus on macrophages and broadly dampen induction of the inflammasome.

Autopsies of patients succumbing to severe COVID-19 have linked disease pathology to the accumulation of aberrantly activated macrophages in lungs^59^ and hyperactivated Siglec1+ macrophages co-localizing with SARS-CoV-2 in the hilar LNs of autopsy samples^60^. Grant et al proposed a model in which alveolar macrophages act as “Trojan horses” that traffic to adjacent lung regions, ferrying cell-associated virus and propagating inflammation^61^. In this context, the observation that IFNmod was able to potently reduce the expression of the viral attachment receptor Siglec-1 on CD14+ blood monocytes and BAL macrophages, with concomitantly lowered ISG expression and inflammasome activation, provides a possible mechanism by which the IFN system may enhance trans-infection and enable propagation and dissemination of inflamed macrophages in the airspace. Our scRNA-Seq data are consistent with the “Trojan horse” hypothesis and extend it by identifying that the CD163^+^MRC1^-^ and CD163^+^TREM2^+^ populations are the predominant populations in the lung in which the inflammasome is induced.

This study, using an intervention targeting both IFN-α and IFN-β pathways, shows that excessive inflammation driven by type 1 IFN critically contributes to SARS-CoV-2 pathogenesis in RMs, and demonstrates the potential of IFNmod to limit viral replication, SARS-CoV-2 induced inflammation, and COVID-19 severity.

## Methods

### Study Approval

EPC’s animal care facilities are accredited by both the U.S. Department of Agriculture (USDA) and by the Association for Assessment and Accreditation of Laboratory Animal Care (AAALAC). All animal procedures were performed in line with institutional regulations and guidelines set forth by the NIH’s Guide for the Care and Use of Laboratory Animals, 8^th^ edition, and were conducted under anesthesia with appropriate follow-up pain management to minimize animal suffering. All animal experimentation was reviewed and approved by Emory University’s Institutional Animal Care and Use Committee (IACUC) under permit PROTO202100003.

### Animal models

In the uninfected animal study, 4 (2 females and 2 males; average age of 9 years and 2 months) specific-pathogen free (SPF) Indian-origin rhesus macaques (RM; *Macaca mulatta;* **Supplementary Table 1)** were housed at Emory National Primate Research Center (ENPRC) in the BSL-2 facility. All four uninfected RMs were administered 1 mg/day of IFN-I modulator (IFNmod) for four consecutive days. IFNmod was supplied in solution and diluted with PBS and administered intramuscularly in the thigh. Peripheral blood (PB) and BAL collections were performed at pre- and post-administration timepoints as annotated **(Supplementary Fig. 1a).**

In the SARS-CoV-2-infected animal study, 20 (8 females and 12 males; average age of 11 years and 8 months) SPF Indian-origin rhesus macaques (**Supplementary Table 1**) were housed at the ENPRC as previously described^62^ in the ABSL3 facility. RMs were infected with 1.1×10^6^ plaque forming units (PFU) SARS-CoV-2 via both the intranasal (1 mL) and intratracheal (1 mL) routes concurrently. Two SARS-CoV-2-infected animals were excluded due to error in specimen processing, resulting in missing time points. Nine RMs were administered 1 mg/day of IFN-I modulator (IFNmod) starting one day prior to infection (−1 dpi) until 2 dpi. IFNmod was supplied in solution and diluted with PBS and administered intramuscularly in the thigh. The other 9 animals served as untreated, SARS-CoV-2 infected group. At each cageside access RMs were clinically scored for responsiveness, discharges, respiratory rate, respiratory effort, cough, and fecal consistency **(Supplementary Table 2)**. Additionally, at each anesthetic access, body weight, body condition score, respiratory rate, pulse oximetry, and rectal temperature was recorded and RMs were clinically scored for discharges, respiratory character, and hydration **(Supplementary Table 3)**. Longitudinal tissue collections of peripheral blood (PB); bronchoalveolar lavage (BAL); and nasal, and pharyngeal mucosal swabs in addition to thoracic X-rays (ventrodorsal and right lateral views) were performed immediately following IFNmod administration as annotated (**Fig. 1a**). In addition to the tissues listed above, at necropsy the following tissues were processed for mononuclear cells: hilar LN, caudal (lower) lung, cranial (upper) lung, and spleen. Additional necropsy tissues harvested for histology included nasopharynx.

### IFNmod production

IFNmod (also named IFN-1ant) is an IFNα2 engineered protein, including the R120E mutation and the terminal 5 amino acids of IFNα8. This results in tight binding to IFNAR2, and bellow detection binding to IFNAR1^31, 63^. For expression, IFNmod was cloned in a pET28bdSUMO plasmid^64^ and grown in BL21 DE3 cells to OD_600_ of 0.6 at 37°C in 2TY broth, which after cells were transferred to 15°C and 0.2 mM fresh IPTG was added for ON growth. Cells were harvested and disintegrated in 50 mM Tris, 100 mM NaCl buffer (pH 8) including Benzonase (Sigma-Aldrich, Cat#: E1014-5KU), protease inhibitor cocktail (Sigma-Aldrich, Cat#: P8465-5ML) and lysozyme (Sigma-Aldrich, Cat# 62970-5G-F), and purified by NiNTA beads (Ni-NTA beads, Merck, cat. 70666-4). IFNmod fused with SUMO was purified by the on-column cleavage using 50 mM Tris pH 8, 100 mM NaCl and BD-Sumo protease (1:200 from 1mg\ml stock, in-house production) for 2 hr at RT, and then moved to over-night cleavage at 4°C^64^. After elution from NiNTA, IFNmod was subjected to size exclusion chromatography using a Superdex 75 26/60 column. For endotoxin removal, IFNmod was subjected to ToxinEraser^TM^ (GenScript, L00338), with endotoxin levels measured using ToxinSensor^TM^ (GenScript, Cat# l00350). Yields were ∼15 mg pure protein per liter.

### ISGs induction in IFNmod-treated Calu-3 cells

Calu-3 cells were seeded in 48 well plates. 24 hours post-seeding, cells were treated with 0, 0.004, 0.04, or 0.4 μg/ml IFNmod with or without 0.04 µg/ml IFNα. 24 hours later, cells were harvested for qRT-PCR analysis. Total RNA extraction was performed using the Quick-RNA Microprep Kit (Zymo research) according to the manufacturer’s instructions. On a StepOnePlus Real-Time PCR System (Applied Biosystems), reverse transcription and qRT–PCR were performed in one step using a SuperScript III Platinum Kit (Thermo Fisher Scientific) following the manufacturer’s instructions. Each TaqMan probe (OAS1, Hs05048921_s1; ISG15, Hs01921425_s1; CXCL10, Hs00171042_m1; β-Actin, Hs99999903_m1) was acquired as premixed TaqMan Gene Expression Assays (Thermo Fisher Scientific) and added to the reaction. Expression levels of each target gene were calculated by normalizing to Actin levels using the ΔΔCT method.

### Impact of IFNmod on SARS-CoV-2 replication in Calu-3 cells

30,000 Calu-3 cells were seeded in 96 well plates. 24 hours post-seeding cells were treated with 0, 0.004, 0.04, or 0.4 ug/ml IFNmod. 24 hours later, cells were infected with SARS CoV-2 NL-02-2020 (MOI 0.1). 5h post infection cells were washed once with PBS, supplemented with fresh media, and treated again with IFNmod. 48h later, supernatants and cells were harvested for qRT-PCR analysis. N (nucleoprotein) transcript levels were determined in supernatants collected from SARS-CoV-2 infected Calu-3 cells. Total RNA was isolated using the Viral RNA Mini Kit (Qiagen, Cat#52906) according to the manufacturer’s instructions. RT-qPCR was performed as previously described using TaqMan Fast Virus 1-Step Master Mix (Thermo Fisher, Cat#4444436) and a OneStepPlus Real-Time PCR System (96-well format, fast mode). Primers were purchased from Biomers (Ulm, Germany) and dissolved in RNAse free water. Synthetic SARS-CoV-2-RNA (Twist Bioscience, Cat#102024) or RNA isolated from BetaCoV/France/IDF0372/2020 viral stocks quantified via this synthetic RNA (for low CT samples) were used as a quantitative standard to obtain viral copy numbers. All reactions were run in duplicates. Forward primer (HKU-NF): 5’-TAA TCA GAC AAG GAA CTG ATT A-3’; Reverse primer (HKU-NR): 5’-CGA AGG TGT GAC TTC CAT G-3’; Probe (HKU-NP): 5’-FAM-GCA AAT TGT GCA ATT TGC GG-TAMRA-3’.

### Viral Stocks

SARS-CoV-2 (USA-WA/2020 strain) was obtained from BEI Resources (Cat no. NR-53899, Lot: 70040383). Prior to infection, stocks were titered on Vero E6 cells by plaque assay (ATCC, CRL-1586) and sequenced to verify the stock’s genomic integrity.

### Determination of viral load RNA

SARS-CoV-2 genomic RNA was quantified in nasopharyngeal (NP) swabs, throat swabs, and bronchoalveolar lavages (BAL). Swabs were placed in 2mL of PBS (CORNING). Virus was inactivated by mixing 1:1 with Buffer ATL (Qiagen) prior to viral RNA extraction from NP swabs, throat swabs, and BAL on fresh, inactivated specimens. Viral RNA was extracted on the QIASymphony SP platform using the DSP virus/pathogen kit according the manufacturer’s protocols. Quantitative PCR (qPCR) was performed on viral RNA samples using the N2 primer and probe set designed by the CDC for their diagnostic algorithm: CoV2-N2-F: 5’-TTACAAACATTGGCCGCAAA-3’, CoV2-N2-R: 5’-GCGCGACATTCCGAAGAA-3’, and CoV2-N2-Pr: 5’-FAM-ACAATTTGCCCCCAGCGCTTCAG-BHQ-3’. The primer and probe sequences for the sub-genomic mRNA or sgRNA transcript of the E gene are SGMRNA-E-F: 5’-CGATCTCTTGTAGATCTGTTCTC-3’, SGMRNA-E-R: 5’-ATATTGCAGCAGTACGCACACA-3’, and SGMRNA-E-Pr: 5’-FAM-ACACTAGCCATCCTTACTGCGCTTCG-3’. qPCR reactions were performed in duplicate with the TaqMan Fast Virus 1-step Master Mix using the manufacturer’s cycling conditions, 200nM of each primer, and 125nM of the probe. The limit of detection in this assay was 70 copies per mL of PBS or BAL. To verify sample quality the CDC RNase P p30 subunit qPCR was modified to account for rhesus macaque specific polymorphisms. The primer and probe sequences are RM-RPP30-F 5’-AGACTTGGACGTGCGAGCG-3’, RM-RPP30-R 5’-GAGCCGCTGTCTCCACAAGT-3’, and RPP30-Pr 5’-FAM-TTCTGACCTGAAGGCTCTGCGCG-BHQ1-3’. A single well from each extraction was run as above to verify RNA integrity and sample quality via detectable and consistent cycle threshold values. All quantities were determined using *in vitro* transcribed RNA standards that were verified with quantified viral RNA stocks acquired from ATCC and BEI Resources. Using remaining viral RNA extracted from NP swabs and BAL, sgRNA E mRNA viral loads were repeated by the NIAD Vaccine Research Center (VRC) as described ^65, 66^ using the following primer and probe sets: sgLeadSARSCoV2_F (5′-CGATCTCTTGTAGATCTGTTCTC-3′), E_Sarbeco_P (5′-FAM-ACACTAGCCATCCTTACTGCGCTTCG-BHQ1-3′), and E_Sarbeco_R (5′-ATATTGCAGCAGTACGCACACA-3′). Viral sgRNA N mRNA was also quantified by the VRC using the following primer and probe sets: Forward: 5′-CGATCTCTTGTAGATCTGTTCTC-3′, Probe: 5′-FAM-TAACCAGAATGGAGAACGCAGTGGG-BHQ1–3′, Reverse: 5′-GGTGAACCAAGACGCAGTAT-3′). The lower limit of detection for the assay conducted at the VRC was 50 copies per mL of PBS or BAL.

### SARS-CoV-2 quantification from necropsy samples

Briefly, an approximately 0.5 cm^3^ sample of each tissue was collected at necropsy, placed in 1mL RNAlater Solution (invitrogen), stored at 4°C for 24 hours, and then frozen at −80°C. ∼50mg of each thawed sample was transferred to a microcentrifuge tube containing a 5mm stainless steel bead (Qiagen, Part No. 69989) and 1200 µL Buffer RLT Plus, and homogenized twice for 1-5 minutes each at 25Hz using a TissueLyser LT sample disruptor (Qiagen, Part No. 85600) with a 12 tube Tissue Lyser LT adapter (Qiagen, Part No. 69980). Following centrifugation of the samples at maximum speed for 3 minutes, the supernatant of the samples was collected. Tissue RNA was extracted from homogenized supernatants on the Qiasymphony SP platform using the QIAsymphony RNA Kit (Qiagen, Part No. 931636). Viral RNA was quantified as above and normalized to the copy number of RPP30.

### Histopathology and immunohistochemistry

Due to study end point, the animals were euthanized, and a complete necropsy was performed. For histopathologic examination, various tissue samples including lung, nasal turbinates, trachea, or brain, were fixed in 4% neutral-buffered paraformaldehyde for 24h at room temperature, routinely processed, paraffin-embedded, sectioned at 4μm, and stained with hematoxylin and eosin (H& E). The H&E slides from all tissues were examined by two board certified veterinary pathologists. For each animal, all the lung lobes were used for analysis and affected microscopic fields were scored semi-quantitatively as Grade 0 (None); Grade 1 (Mild); Grade 2 (Moderate) and Grade 3 (Severe). Scoring was performed based on these criteria: number of lung lobes affected, type 2 pneumocyte hyperplasia, alveolar septal thickening, fibrosis, perivascular cuffing, peribronchiolar hyperplasia, inflammatory infiltrates, hyaline membrane formation. An average lung lobe score was calculated by combining scores from each criterion. Digital images of H&E stained slides were captured at 40× and 200× magnification with an Olympus BX43 microscope equipped with a digital camera (DP27, Olympus) using Cellsens® Standard 2.3 digital imaging software (Olympus).

Immunohistochemical (IHC) staining of sections of lung was performed using a biotin-free polymer system. The paraffin-embedded sections were subjected to deparaffinization in xylene, rehydration in graded series of ethanol, and rinsed with double distilled water. Antigen retrieval was performed by immersing sections in DIVA Decloaker (Biocare Medical) at 125 LJC for 30 seconds in a steam pressure decloaking chamber (Biocare Medical) followed by blocking with Background Sniper Reagent (Biocare Medical) for 10 minutes. The sections were incubated with Thyroid Transcription Factor-1 (Clone 8G7G3/1) for overnight at 4°C followed by a detection polymer system (MACH 2™; Biocare Medical). Labeled antibody was visualized by development of the chromogen (DAB Chromogen Kits; Biocare Medical).

Tissues were fixed in freshly prepared 4% paraformaldehyde for 24 h, transferred to 70% ethanol, paraffin embedded within 7-10 days, and blocks sectioned at 5 µm. Slides were baked for 30-60 min at 65°C then deparaffinized in xylene and rehydrated through a series of graded ethanol to distilled water. Heat induced epitope retrieval (HIER) was performed with the antigen retrieval buffers citraconic anhydride (0.01% with 0.05% Tween; Mx1, Iba-1, and Ki-67) or citrate buffer (pH 6.0; MPO) in a Biocare NxGen Decloaking Chamber that was set to 110°C for 15 min. The slides were cooled, rinsed twice in distilled water and 1X TBS with 0.05% Tween-20 (TBS-T), blocked (TBS-T + 0.25% casein) for 30 minutes at room temperature, then incubated at room temperature with antibodies against Mx1 (EMD; Cat. No. MABF938 at 1:1000 for 1 hour), MPO (Dako; Cat. No. A0398 at 1:1000 for 1 hour), Iba-1 (BioCare; Cat. No. CP290A at 1:500 for 1 hour), and Ki67 (BD Pharmingen; Cat. No. 550609 at 1:200 for 1 hour). Endogenous peroxidases were blocked with 1.5% H_2_O_2_ in TBS-T for 10 minutes. Slides were then incubated with Rabbit Polink-1 HRP (GBI Labs; Cat. No. D13-110 for MPO and Iba-1) and Mouse Polink-2 HRP (GBI Labs; Cat. No. D37-110 for Mx1 and Ki67). Slides were developed using Impact™ DAB (3,3′-diaminobenzidine; Vector Laboratories), washed in ddH_2_O, counterstained with hematoxylin, mounted in Permount (Fisher Scientific), and scanned at 20x magnification on an Aperio AT2 (Leica Biosystems). Staining for MPO, Mx1, Iba-1, and Ki67 IHC was performed as previously described using a Biocare intelliPATH autostainer.

### Tissue Processing

PB was collected from the femoral vein in sodium citrate, serum separation, and EDTA tubes from which plasma or serum was separated by centrifugation within 1 hour of phlebotomy. Serum from blood collected in serum separator tubes and plasma from blood collected in sodium citrate tubes were used for comprehensive blood chemistry panels. EDTA PB was used for complete blood counts, whole blood staining, and PBMC isolation and staining. Following the centrifugation of EDTA PB and the removal of plasma, 500uL of the remaining fraction of blood was lysed with ACK lysis buffer, pelleted via centrifugation, and washed twice with PBS in preparation for whole blood flow cytometry staining. Peripheral blood mononuclear cells (PBMCs) were also isolated from the fraction of EDTA blood remaining following the removal of plasma using a Ficoll-Paque Premium density gradient (GE Healthcare), and washed with R-10 media. R-10 media was composed of RPMI 1640 (Corning) supplemented with 10% heat-inactivated fetal bovine serum (FBS), 100 IU/mL penicillin, 100 μg/mL streptomycin, and 200 mM L-glutamine (GeminiBio).

Nasopharyngeal swabs were collected under anesthesia by using a clean flocked nylon-tipped swab (Azer Scientific Inc, Nasopharyngeal swab Individually wrapped, sterile, 76422-824, VWR International) placed approximately 2-3cm into the nares. Oropharyngeal swabs were collected under anesthesia using polyester tipped swabs (Puritan Standard Polyester Tipped applicator, polystyrene handle, 25-806 2PD, VWR International) to streak the tonsils and back of throat bilaterally (throat/pharyngeal). The swabs were dipped in 2 mL PBS (Corning) and vortexed for 30 sec, and the eluate was collected.

To collect BAL, a fiberoptic bronchoscope (Olympus BF-XP190 EVIS EXERA III ULTRA SLM BRNCH and BF-P190 EVIS EXERA 4.1mm) was manipulated into the trachea, directed into the primary bronchus, and secured into a distal subsegmental bronchus upon which 35-70 mL of normal saline (0.9% NaCl) was administered into the bronchus and re-aspirated to obtain a minimum of 20ml of lavage fluid. At necropsy, a large sterile syringe with the plunger removed was inserted into the tracheal opening and approximately 150 ml of sterile 1 x PBS was poured into both sides of the lungs. The syringe was then removed and re-inserted into the tracheal opening with the plunger replaced. Lavage fluid was then pulled back into syringe and dispensed into conical tubes. BAL was filtered through a 70μm cell strainer and multiple aliquots were collected for viral loads. Next, the remaining BAL was centrifuged at 2200rpm for 5 minutes and the BAL fluid supernatant was collected for mesoscale analysis. Pelleted BAL cells were resuspended in R10 and used for downstream analyses including 10x, Bulk RNAseq, and flow cytometry.

The lower portions of the cranial (upper) lung lobes and the bilateral upper portions of the caudal (lower) lung lobes were collected at necropsy. Lung tissue was injected with digestion media containing 0.4mg/mL DNase I (StemCell Technologies), 2.5 mg/mL of Collagenase D (Roche), and 0.2mg/mL Liberase TL Research Grade (Sigma-Aldrich) in HBSS using a blunt end needle. Next, lung tissue was cut into small pieces using blunt end scissors and incubated in digestion media at 37C for 1 hour. Lung tissue was then transferred into gentleMACS C tubes (Miltenyi Biotec) and homogenized using a gentleMACS Dissociator, program “Lung 02_01” (Miltenyi Biotec). The resulting homogenized tissue was filtered through a 100μm BD Falcon cell strainer and washed with R-10 media.

Hilar LN biopsies were collected at necropsy, sectioned using blunt, micro-dissection scissors and mechanically disrupted through a 70μm cell strainer and washed with R-10 media.

Mononuclear cells were counted for viability using a Countess II Automated Cell Counter (Thermo Fisher) with trypan blue stain and were cryo-preserved in aliquots of up to 2×10^7^ cells in 10% DMSO in heat-inactivated FBS. Whole tissue segments (0.5 cm^3^) were snap frozen dry or stored in RNAlater (Qiagen) for analyses of RNA-seq and tissue viral quantification, respectively.

### Bulk and single-cell RNA-Seq Library and sequencing from NHP BALs

Single cell suspensions from BAL were prepared in a BSL3 as described above for flow cytometry. For bulk RNA-Seq, 50,000 cells were lysed directly into 700 ul of QIAzol reagent and RNA was isolated using the miRNeasy Micro kit (Qiagen) with on-column DNase digestion. RNA quality was assessed using an Agilent Fragment Analyzer and ten nanograms of total RNA was used as input for cDNA synthesis using the Clontech SMART-Seq v4 Ultra Low Input RNA kit (Takara Bio) according to the manufacturer’s instructions. Amplified cDNA was fragmented and appended with dual-indexed bar codes using the NexteraXT DNA Library Preparation kit (Illumina). Libraries were validated by capillary electrophoresis on an Agilent Fragment Analyzer, pooled at equimolar concentrations, and sequenced on an Illumina NovaSeq6000. For single-cell RNA-Seq, approximately 20,000 cells were loaded onto the 10X Genomics Chromium Controller in the BSL3 facility using the Chromium NextGEM Single Cell 5’ Library & Gel Bead kit according to manufacturer instructions^67^. The resulting gene expression libraries were sequenced as paired-end 26×91 reads on an Illumina NovaSeq6000 targeting a depth of 50,000 reads per cell. Cell Ranger software was used to perform demultiplexing of cellular transcript data, and mapping and annotation of UMIs and transcripts for downstream data analysis.

### Bulk RNA-Seq analysis

Reads were aligned using STAR v2.7.3.^68^. The STAR index was built by combining genome sequences for Macaca mulatta (Mmul10 Ensembl release 100), SARS-CoV2 (strain MT246667.1 - NCBI). The gffread utility (https://github.com/gpertea/gffread) was used to convert gff3 file for SARS-CoV2 and the resulting gtf file for SARS-CoV2 was edited to include exon entries which had the same coordinates as CDS to get counts with STAR. The combined genomic and gtf files were used for generating the STAR index. Transcript abundance estimates were calculated internal to the STAR aligner using the algorithm of htseq-count^69^. The ReadsPerGene files were read in R constructing a count matrix that was imported in DESeq2 using the DESeqDataSetFromMatrix function. DESeq2 was used for normalization^70^, producing both a normalized read count table and a regularized log expression table. Only the protein coding genes defined in the gtf file were used for analysis. The design used was: ∼ Subject + Treatment * Timepoint. The regularized log expression values were obtained using the rlog function with the parameters blind =FALSE and filtType = “parametric”. GSEA 4.1.0 (https://www.gsea-msigdb.org/) was used for gene set enrichment analysis with the following gene sets: Hallmark and Reactome and Biocarta Canonical pathways (MsigDB), NHP ISGs^34^. GSEA was run with default parameters with the permutation type set to gene_set. The input for GSEA was the regularized log expression values obtained from DESeq2 which was filtered to remove genes with mean expression <=0. The regularized log expression values were also used to generate heatmaps using the Complex Heatmap R library.

### Bulk RNA-Seq Whole blood analysis

Whole blood was collected from RM into PAXgene RNA tubes (PreAnalytiX) according to the manufacturer’s instructions. RNA was isolated using RNAdvance Blood Kit (Beckman) following the manufacturer’s instructions. mRNAseq libraries were constructed using KAPA RNA HyperPrep Kit with RiboErase (HMR) Globin in conjunction with the KAPA mRNA Capture Kit (Roche Sequencing). Libraries were sequenced on an Illumina NextSeq500 sequencer using Illumina NextSeq 500/550 High Output v23 (150 cycles) following the manufacturer’s protocol for sample handling and loading. Reads were initially filtered for hemoglobin and rRNA reads using bowtie 2.4.2, and then aligned to the *Macaca mulatta* (Mmul10 Ensembl release 100) genome using STAR v2.7.5a. Final counts were then generated with HTSeq-count v0.13^68, 69, 71^. Genes with a mean count across samples lower than 5 were filtered from analysis. Counts were normalized using TMM normalization/voom^72–74^. Differential gene analysis was done using limma with design (∼0 +time:treatment + animal) with an adjusted pval cutoff of 0.05 and absolute LFC value greater than 0.58 comparing 1,2,4,5 and 7 dpi to baseline at each time-point for animals treated with IFNmod and untreated animals (infected with SARS-CoV2)^74^. Over representation analysis was performed on significant DE genes using SetRank^75^. To identify pathways significantly enriched (adjusted P val < 0.05). Activity of pathways of interest were illustrated by calculating the mean logCPM value of genes in each pathway across time.

### Single-cell RNA-Seq Bioinformatic Analysis

The cellranger v6.1.0 (10X Genomics) pipeline was used for processing the 10X sequencing data and the downstream analysis was performed using the Seurat v4.0.4^76^ R package. A composite reference comprising of Mmul10 from Ensembl release 100 and SARS-CoV2 (strain MT246667.1 - NCBI) was used for alignment with cellranger. The percentage of SARS-CoV2 reads was determined using the PercentageFeatureSet for SARS-CoV2 genes.

### BAL samples

The samples were demultiplexed using HTODemux function in Seurat. The gene expression matrix was filtered to include protein coding genes and exclude genes encoded on Y chromosome, B and T cell receptor genes, mitochondrial genes, RPS and RPL genes and SARS-CoV2 genes. The cells were further filtered on the following criteria: nFeature_RNA >=500 and <= 3500, ncount_RNA >=250 and log10GenesPerUMI > 0.8. After filtering, the samples were normalized using SCTransform method^77^ and integrated using the first 30 dimensions with the default CCA method ^78^. Three samples were dropped - two due to low cell numbers (RNi17 2 dpi and RZs14 7 dpi) and one for large proportion of epithelial cells (CD68 baseline). The downstream analyses was carried out with the remaining samples comprising of 62,081cells.

The integrated object was split into individual samples and after filtering the three samples, the remaining samples were normalized using the SCTransform method^77^ and then integrated using the reciprocal PCA method ^79^. The first 30 dimensions were used with the FindIntegrationAnchors, FindUMAP and FindNeighbors method. Clustering was carried out using the default Louvain method and the resolution was set to 1. Cell annotations were carried out based on the expression of canonical markers in seurat clusters and SingleR v1.4.0 library (Blueprint Encode database) ^80^ annotations were used as a guide. As a distinct cluster could not be determined for neutrophils based on the expression of canonical marker genes, the SingleR annotations were used for neutrophils. Differential gene expression analysis was carried out using the FindMarkers function with “MAST”^81^ method.

To further classify the macrophages/monocytes in BAL, only cells in the largest cluster comprising the macrophages/monocytes were further processed. The subset function was used to get these cells followed by splitting the object in individual samples. The MapQuery function was used to map individual samples to a reference lung macrophage/monocyte dataset from three healthy rhesus macaques obtained from a previously published study^82^ using the same approach that we have previously described^52^.

### Lung samples

We processed lung samples from three animals in each group at 2 dpi and one each from 7 dpi. Two of the 2 dpi samples were dropped (one from each group), due to low cell recovery. The cellranger pipeline was used as described above. The filtered counts were read into Seurat using the Read10X_h5 function. The gene expression matrix was filtered to include protein coding genes and exclude genes encoded on Y chromosome, B and T cell receptor genes, mitochondrial genes, RPS and RPL genes and SARS-CoV2 genes. The cells were filtered based on the following criteria: (i) nFeature_RNA between 500 and 5000 and (ii) percent mitochondrial genes <= 25%. Normalization was performed using SCTransform and the samples were integrated using reciprocal PCA. The first 30 dimensions were used and clustering was carried out with the resolution set to 0.1 using the default Louvain algorithm in seurat. The clusters were annotated based on the expression of canonical markers and roughly divided into four major subsets: epithelial, myeloid, lymphoid and others. Each subset was then clustered separately to fine tune the cell type annotations. The human Lung v1 reference^54^ in Azimuth (https://azimuth.hubmapconsortium.org/) was used to guide the cell annotations. Based on the expression of canonical markers, some clusters were classified as doublets and some remained unassigned. After removing the doublets and unassigned clusters, UMAPs showed some additional cells that coincided with the removed doublets/unassigned clusters and these were removed as well. Finally, a total of 42,699cells were used subsequently for downstream analysis. Differential gene expression analysis was carried out using the FindMarkers function with “MAST” method. Over –representation analysis was carried out using clusterProfiler v4.5.0.992^83^ with Hallmark, Reactome, KEGG and BioCarta genesets from the msigdb database^84–86^. The msigdbr v7.5.1 library (https://igordot.github.io/msigdbr/) was used for retrieving the msigdb databases.

### Immunophenotyping

23-parameter flow cytometric analysis was perform on fresh PBMCs and mononuclear cells (10^6^ cells) derived from LN biopsies, BAL, and lung. Immunophenotyping was performed using anti-human monoclonal antibodies (mAbs), which we^62, 87–89^ and others, including databases maintained by the NHP Reagent Resource (MassBiologics), have shown as being cross-reactive in RMs. A panel of the following mAbs was used for longitudinal T-cell phenotyping in PBMCs: anti-CCR7-BB700 (clone 3D12; 2.5 μL; cat. # 566437), anti-CXCR3-BV421 (clone IC6; 2.5 μL; cat. # 562558), anti-Ki-67-BV480 (clone B56; 5 μL; cat. # 566109), anti-CCR4-BV750 (clone 1G1 1E5; 2.5 μL; cat. # 746980), anti-CD3-BUV395 (clone SP34-2; 2.5 μL; cat. # 564117), anti-CD8-BUV496 (clone RPA-T8; 2.5 μL; cat. # 612942), anti-CD45-BUV563 (clone D058-1283; 2.5 μL; cat. # 741414), anti-CD49a-BUV661 (clone SR84; 2.5 μL; cat. # 750628), anti-CD28-BUV737 (clone CD28.2; 5 μL; cat. # 612815), anti-CD69-BUV805 (clone FN50; 2.5 μL; cat. # 748763), and Fixable Viability Stain 700 (2 μL; cat. # 564997) all from BD Biosciences; anti-CD95-BV605 (clone DX2; 5 μL; cat. # 305628), anti-HLA-DR-BV650 (clone L243; 5 μL; cat. # 307650), anti-CD25-BV711 (clone BC96; 5 μL; cat. # 302636), anti-PD-1-BV785 (clone EH12.2H7; 5 μL; cat. # 329930), anti-CD101-PE-Cy7 (clone BB27; 2.5 μL; cat. # 331014), anti-FoxP3-AF647 (clone 150D; 5 μL; cat. # 320014), and anti-CD4-APC-Cy7 (clone OKT4; 2.5 μL; cat. # 317418) all from Biolegend; anti-CD38-FITC (clone AT1; 5 μL; cat. # 60131FI) from STEMCELL Technologies; and anti-CXCR5-PE (clone MU5UBEE; 5 μL; cat. # 12-9185-42), anti-GranzymeB-PE-TexasRed (clone GB11; 2.5 μL; cat. # GRB17), and anti-CD127-PE-Cy5 (clone eBioRDR5; 5 μL; cat. # 15-1278-42) all from Thermo Fisher. mAbs for chemokine receptors (i.e. CCR7) were incubate at 37°C for 15 min, and cells were fixed and permeabilized for 30 min at room temperature using a FoxP3 / Transcription Factor Staining Buffer Kit (Tonbo Biosciences; cat. # TNB-0607-KIT). A panel of the following mAbs was used for the longitudinal phenotyping of innate immune cells in whole blood (500 μL), as described in^90^, and mononuclear cells (2×10^6^ cells) derived from LN biopsies, BAL, and lung: anti-CD20-BB700 (clone 2H7; 2.5 μL; cat. # 745889), anti-CD11b-BV421 (clone ICRFF44; 2.5 μL; cat. # 562632), anti-Ki-67-BV480 (clone B56; 5 μL; cat. # 566109), anti-CD14-BV605 (clone M5E2; 2.5 μL; cat. # 564054), anti-CD56-BV711 (clone B159; 2.5 μL; cat. # 740781), anti-CD163-BV750 (clone GHI/61; 2.5 μL; cat. # 747185), anti-CD3-BUV395 (clone SP34-2; 2.5 μL; cat. # 564117), anti-CD8-BUV496 (clone RPA-T8; 2.5 μL; cat. # 612942), anti-CD45-BUV563 (clone D058-1283; 2.5 μL; cat. # 741414), anti-CCR2-BUV661 (clone LS132.1D9; 2.5 μL; cat. # 750472), anti-CD16-BUV737 (clone 3G8; 2.5 μL; cat. # 564434), anti-CD101-BUV805 (clone V7.1; 2.5 μL; cat. # 749163), anti-CD169-PE (clone 7-239; 2.5 μL; cat. # 565248), and anti-CD206-PE-Cy5 (clone 19.2; 20 μL; cat. # 551136) and Fixable Viability Stain 700 (2 μL; cat. # 564997) all from BD Biosciences; anti-ACE2-AF488 (clone Polyclonal; 5 μL; cat. # FAB9332G-100UG) from R & D; anti-HLA-DR-BV650 (clone L243; 5 μL; cat. # 307650), anti-CD11c-BV785 (clone 3.9; 5 μL; cat. # 301644), and anti-CD123-APC-Fire750 (clone 315; 2.5 μL; cat. # 306042) all from Biolegend; anti-GranzymeB-PE-TexasRed (clone GB11; 2.5 μL; cat. # GRB17) from Thermo Fisher; anti-CD66abce-PE-Vio770 (clone TET2; 1 μL; cat. # 130-119-849) from Miltenyi Biotec; anti-NKG2A-APC (clone Z199; 5 μL; cat. # A60797) from Beckman Coulter. mAbs for chemokine receptors (i.e. CCR2) were incubated at 37°C for 15 min, and cells were fixed and permeabilized at room temperature for 15 min with Fixation/Permeabilization Solution Kit (BD Biosciences; cat. #554714). For each sample a minimum of 1.2×10^5^ stopping gate events (live CD3^+^ T-cells) were recorded. All samples were fixed with 4% paraformaldehyde and acquired within 24 hours of fixation. Acquisition of data was performed on a FACSymphony A5 (BD Biosciences) driven by FACS DiVa software and analyzed with FlowJo (version 10.7; Becton, Dickinson, and Company).

### Mesoscale

Plasma was collected from EDTA blood following centrifugation at 2500 rpm for 15 minutes. BALF supernatant was obtained from BAL that was filtered through a 70μm cell strainer and spun down at 1800rpm for 5 minutes. Both plasma and BALF were frozen at −80C and later thawed immediately prior to use.

Cytokines were measured using Mesoscale Discovery. Pro-and anti-inflammatory cytokines and chemokines were measured as part of V-Plex (#K15058D-1). INF alpha was measured with a U plex (K156VHK-1 Mesoscale Discovery, Rockville, Maryland).

Levels for each cytokine/chemokine were determined in double and following the instructions of the kits. The plates were read on a MESO Quick plex 500 SQ120 machine.

Quantities were determined using the discovery Work bench software for PC (version 4.0).

### Quantification and Statistical Analysis

All statistical analyses were performed two-sided with p-values ≤0.05 deemed significant. Ranges of significance were graphically annotated as follows: *, p<0.05; **, p<0.01; ***, p<0.001; ****, p<0.0001. Analyses, unless otherwise noted, were performed with Prism version 8 (GraphPad).

## Resource Availability

### Lead Contact

Further information and requests for resources and reagents should be directed to and will be fulfilled by the Lead Contact, Dr. Mirko Paiardini.

## Materials Availability

This study did not generate new unique reagents.

## Data Availability Statement

Source data supporting this work are available from the corresponding author upon reasonable request. The following sequencing data will be deposited in GenBank: SARS-CoV-2 viral stock (accession # PENDING). Data tables for expression counts for bulk and single-cell RNA-Seq for BAL are deposited in NCBI’s Gene Expression Omnibus and are accessible through the Gene Expression Omnibus (GEO) accession GSE205429. Transcriptomics data sets from whole blood RNA sequencing are deposited at the GEO accession number GSE207665.

## Code Availability Statement

Custom scripts and supporting documentation on the RNA-Seq analyses will be made available at https://github.com/BosingerLab/NHP_COVID_IFNmod. The R markdown code applied to the analyses of whole blood can be accessed at https://github.com/galelab/Paiardini_Modulation_type_I_Interferon.

## Dedication

We would like to dedicate this manuscript to Dr. Timothy N. Hoang, whose commitment to this project and to scientific discovery were crucial in propelling this study forward. Dr. Hoang will be remembered for his intelligence, drive, and love for science and all of the lives that he touched during his short but impactful career.

## Acknowledgments

We thank the Emory National Primate Research Center (EPC) Division of Animal Resources, especially Joyce Cohen, Sherrie M. Jean, and Rachelle L. Stammen in Veterinary Medicine and Stephanie Ehnert, Stacey Weissman, Denise Bonenberger, John M. Wambua, and Racquel Sampson-Harley in Research Resources for providing support in animal care. We would also like to thank Sanjeev Gumber in the EPC Division of Pathology, Kalpana Patel in the EPC Safety Office, Shelly Wang in the Emory CFAR Core, Krista Krish in the EPC Virology Core, Nichole Arnett in the EPC Clinical Pathology Lab, and Elizabeth Beagle, Hadj Aoued, Tristan Horton, David Cowan, Sydney Hamilton and Thomas Hodder in the Emory NPRC Genomics Core. Additionally, we would like to thank Erin Haupt and the Immunologic Core, Department of Immunology, TNPRC for help with MSD analysis, and Elise Smith and Jean Chang in the Gale Lab at WaNPRC for blood RNA-Seq library preparation services. This work was funded by P510D011132-60S4: COVTEN/ACTIV; the Emory MP3 Grant; Fast Grants #2144, and the Pitts Foundation (all to M.Pa). Support for this work was also provided by award NIH Office of Research Infrastructure Programs (ORIP) P51OD11132 to EPC, 1RO1 HL140223 (RDL), and P51 OD010425 (WaNPRC; MG, LW, JT-G). We thank Jana-Romana Fischer and Kerstin Regensburger for excellent technical assistance. F.K. and K.M.J.S. were supported by grants from the German Research Foundation (DFG; CRC 1279 and SP1600/4-1) and the German Federal Ministry for Research and Education (BMBF; Restrict SARS-CoV-2 and IMMUNOMOD). M.H. is part of the International Graduate School of Molecular Medicine, Ulm. Next generation sequencing services and 10X Genomics captures were performed in the Emory NPRC Genomics Core, which is supported in part by NIH P51OD011132. Sequencing data was acquired on an Illumina NovaSeq6000 funded by NIH S10OD026799 to S.E.B. The content of this publication does not necessarily reflect the views or policies of the U.S. Department of Health and Human Services, nor does it imply endorsement of organizations or commercial products.

## Author Contributions

Conceptualization, T.N.H., E.G.V., M.Pi., J.L.H., D.C.D., R.P.J., G.Sc., S.E.B., and M.Pa.; Methodology, T.N.H., E.G.V., A.A.U., G.K.T., M.Pi., R.N., M.H., M.Gag., A.K.B., K.L.P., J.T.-G., L.S.W., M.R., S.K., E.H.C, S.P.K, T.H.V., M.V.; Validation, S.M., M.Gag., and D.C.D.; Formal Analysis, T.N.H., E.G.V., A.A.U., G.K.T., R.N., M.H., J.T.-G., and L.S.W.; Investigation, T.N.H., E.G.V., Z.S., M.Pi., R.N., M.H., M.Gag., K.N., J.T.-G., L.S.W., K.A.K., M.R., S.K., E.H.C., C.W., J.S.W., F.C.-S., R.D.L., M.Gal., K.B.-S., J.E., and M.V.; Resources, S.M. and G.Sc.; Data Curation, A.A.U., A.K.B, K.L.P., J.T.-G, and L.S.W.; Writing – Original Draft, T.N.H., E.G.V., A.A.U., S.E.B., and M.Pa.; Writing – Review & Editing, T.N.H., E.G.V., A.A.U., Z.S., M.Pi, J.L.H., M.Gal., G.Si., K.M.J.S., F.K., G.Sc., S.E.B., and M.Pa.; Visualization, T.N.H., E.G.V., A.A.U., G.K.T, J.T.-G., L.S.W., and S.E.B.; Supervision, M.Gal., S.E.B., M.Pa.; Funding Acquisition, F.K., R.P.J., and M.Pa.

