## Supplementary Figures 1-10 and Supplementary Tables 1-3 for "Modulation of type I interferon responses potently inhibits SARS-CoV-2 replication and inflammation in rhesus macaques"

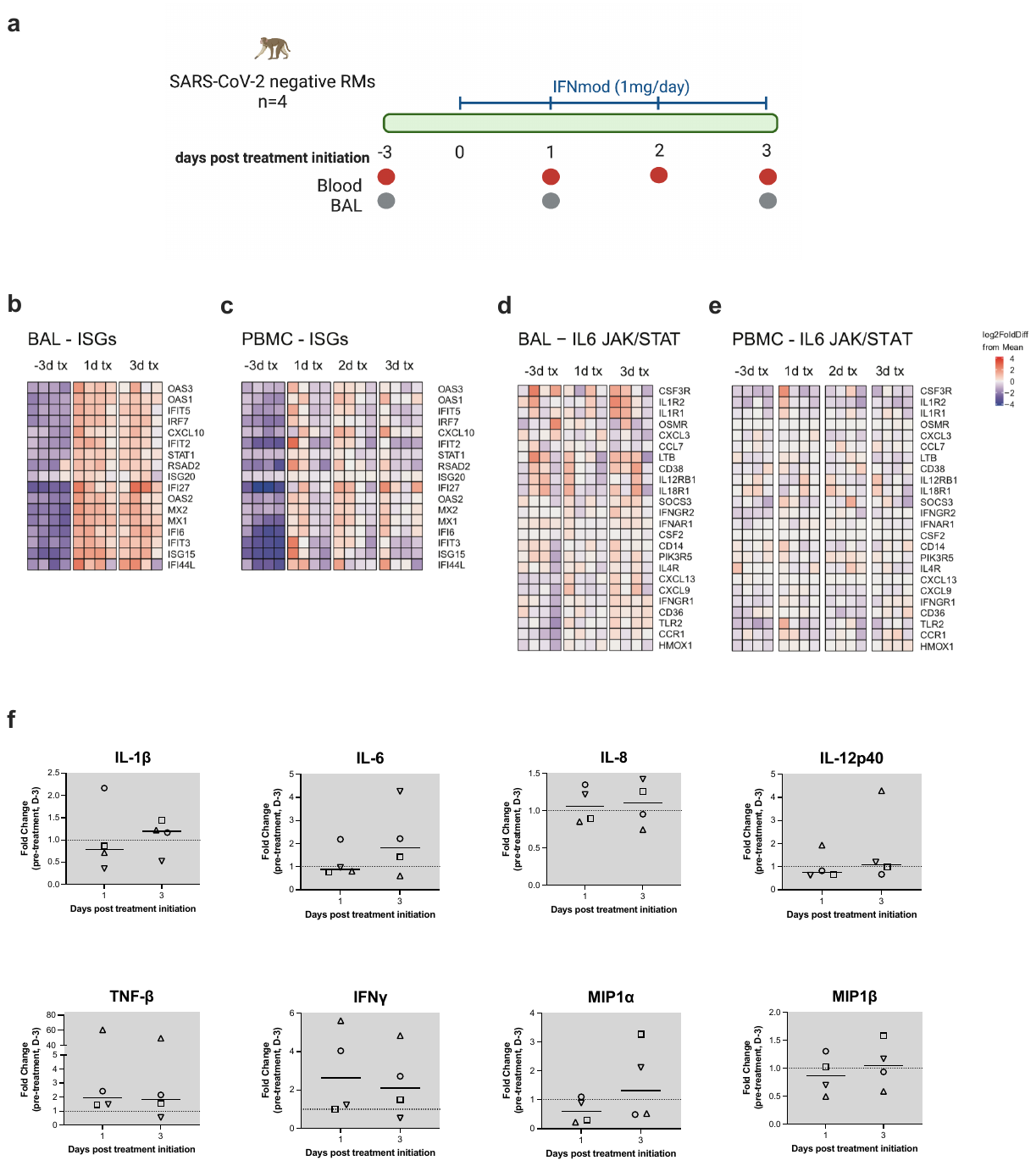

**Supplementary Fig. 1**. **Administration of IFNmod in uninfected RMs resulted in modest upregulation of ISGs without changes to inflammatory genes or inflammatory cytokines and chemokines. (a)** Study design of IFNmod treatment in uninfected RMs; 1mg IFNmod was administered intramuscularly to four uninfected RMs for four consecutive days. Blood and BAL were collected at pre-treatment baseline (3 days before treatment initiation) and once a day from days 1-3 post treatment initiation, with the exception of day 2 post treatment initiation, where only blood was collected. Heatmaps of ISG expression **(b, c)** and expression of genes associated with IL-6 signaling and inflammation **(d, e)** in BAL **(b, d)** and PBMC **(c, e)** of uninfected RMs before and after IFNmod treatment. **(f)** Fold change of cytokines and chemokines in BAL fluid relative to pre-treatment baseline (-3 days post treatment initiation) measured by mesoscale. Statistical analyses for were performed using one-tailed Wilcoxon signed-rank tests. Each black, open symbol represents an uninfected RM. Black lines represent median fold change. Gray-shaded boxes indicate that timepoint occurred during IFNmod treatment.

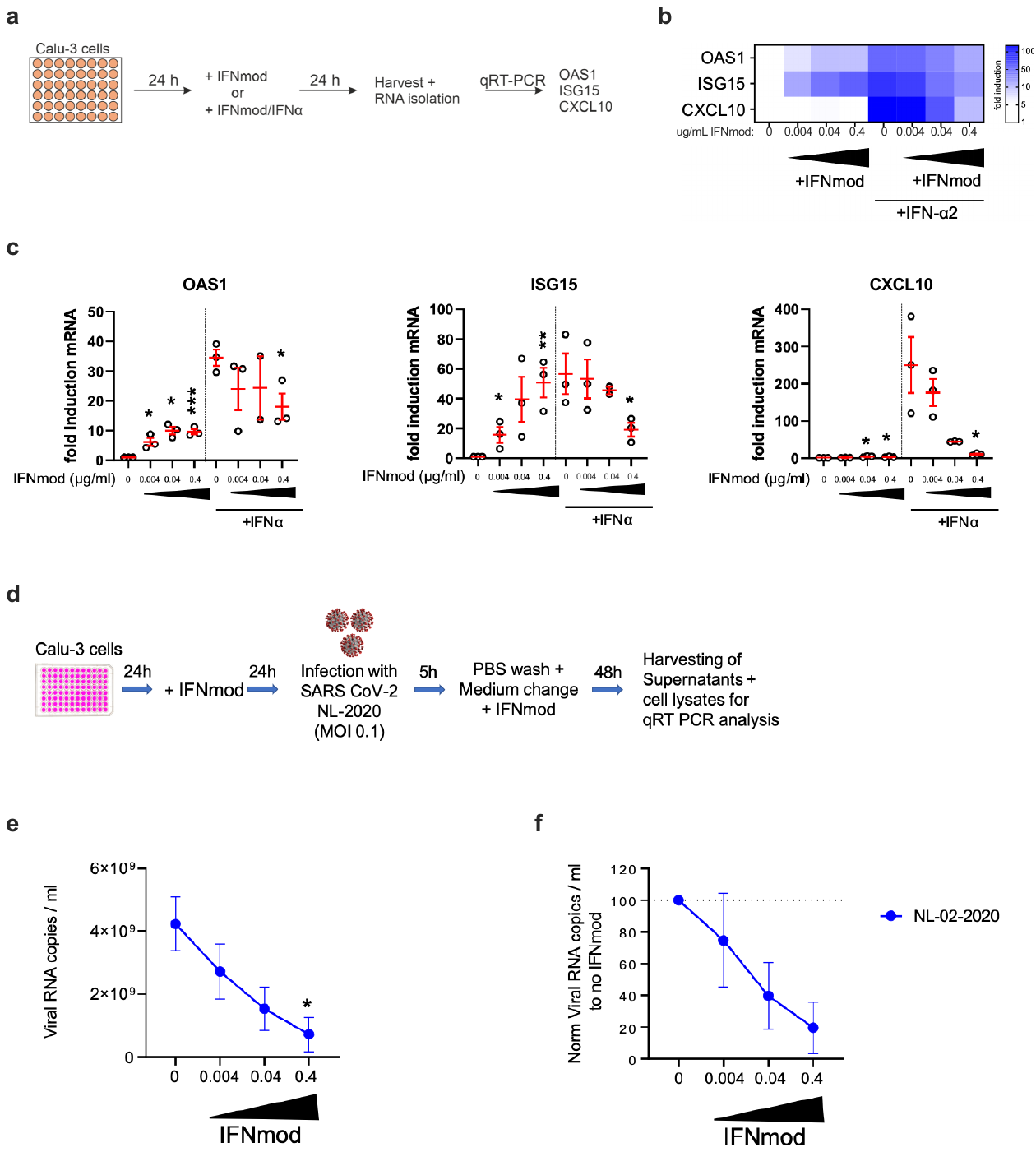

**Supplementary Fig. 2. Administration of IFNmod** **in Calu-3 human primary lung cells modulated type I IFN responses and resulted in inhibition of SARS-CoV-2 replication. (a)** Overview of cell culture setup with IFNmod+/- IFNα treatment. Calu-3 cells were seeded in 48 well plates. 24 hours post seeding, cells were treated with IFNmod (0, 0.0.04, 0.04, or 0.4 µg/ml) with or without IFNα (0.04 µg/ml) as indicated. 24h later, cells were harvested for qRT-PCR analysis. Heatmap **(b)** of and individually plotted **(c)** OAS1, ISG15, and CXCL10 mRNA fold induction in Calu-3 cells following IFNmod (0, 0.0.04, 0.04, or 0.4 µg/ml) +/- IFNα (0.04 µg/ml) treatment from three independent Calu-3 experiments. Statistical analyses were performed using unpaired Student’s t-tests comparing IFNmod +/- IFNα treated samples to their corresponding non-IFNmod treated sample. * p-value < 0.05, ** p-value < 0.01. *** p-value < 0.001. **(d)** Overview of cell culture setup with SARS CoV-2 NL-02-2020 infection. Calu-3 cells were seeded in 96 well plates and treated with IFNmod (0, 0.0.04, 0.04, or 0.4 µg/ml) 24 hours post seeding. 24 hours later, cells were infected with SARS CoV-2 NL-02-2020 (MOI 0.1), 5 hours post-infection cells were washed once with PBS, supplemented with fresh media and treated again with IFNmod. 48 hours later supernatants and cell lysates were harvested for qRT-PCR analysis. **(e)** Viral N RNA copies/mL quantified by qRT-PCR and **(f)** normalized relative to the no IFNmod treatment condition (N= 3 +/- SEM). Statistical analyses for Calu-3 SARS-CoV-2 experiments were performed using unpaired Student’s t-tests comparing IFNmod- treated samples to corresponding non-IFNmod treated samples. * p-value < 0.05, ** p-value < 0.01.

.
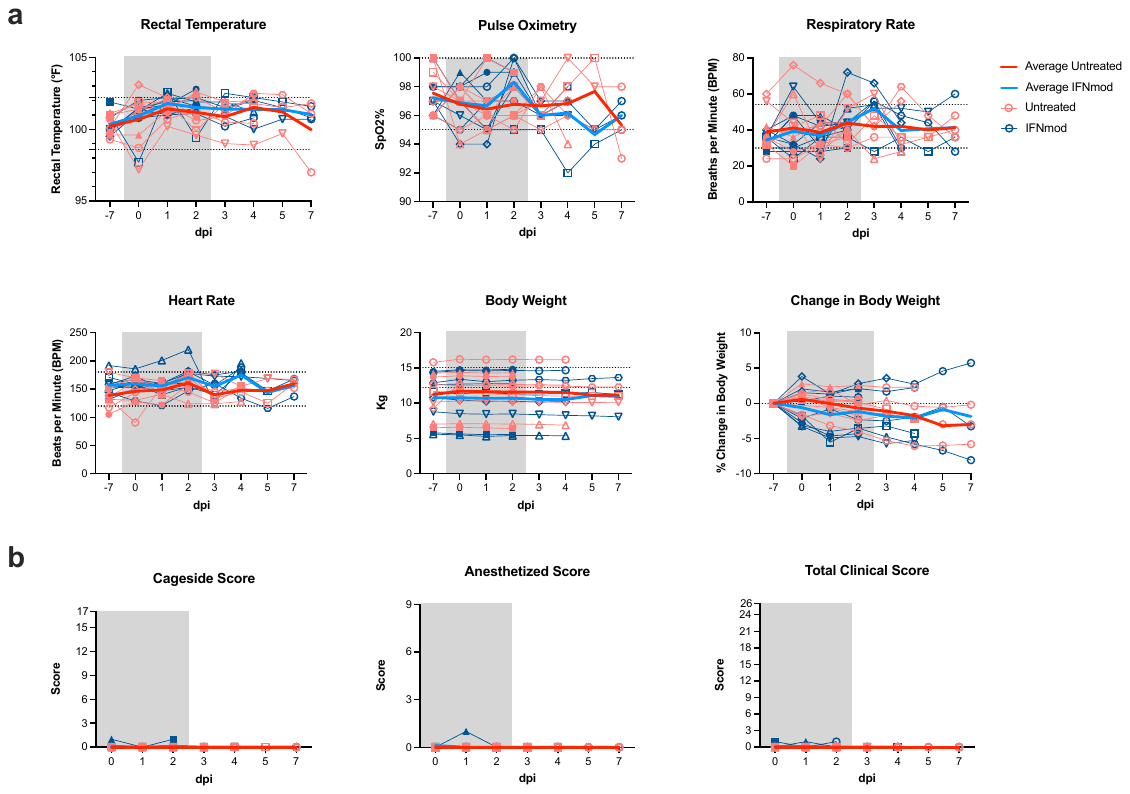

**Supplementary Fig. 3. Administration of IFNmod was safe and well-tolerated in SARS-CoV-2-infected RMs**. **(a)** Longitudinal measurements of rectal temperature, pulse oximetry, respiratory rate, heart rate, body weight, and changes in body weight from pre-infection baseline in untreated (red symbols; n = 9) and IFNmod-treated (blue symbols; n = 9) SARS-CoV-2-infected RMs. **(b)** Cage-side scores, anesthetized scores, and total clinical scores. Black dotted horizontal lines indicate normal ranges for adult indoor RMs. Bolded red and blue lines indicate averages for Untreated and IFNmod treated RMs. Gray-shaded boxes indicate that timepoint occurred during IFNmod treatment.

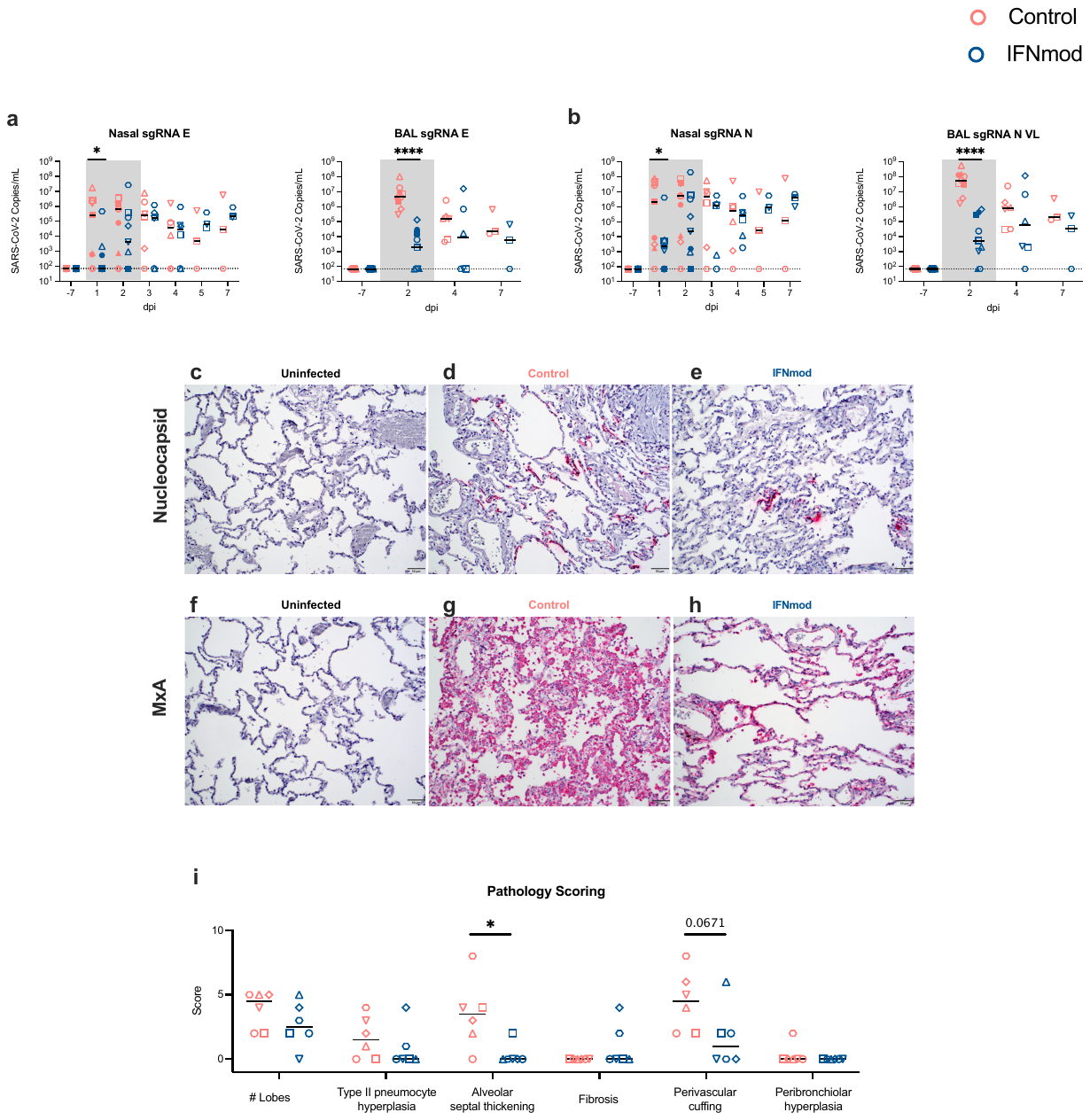
**Supplementary Fig. 4. IFNmod reduced nasopharyngeal and BAL viral loads and nucleocapsid expression, MxA expression, and pathology in lungs in SARS-CoV-2-infected RMs.** sgRNA-E viral loads **(a)** were repeated along with sgRNA-N viral loads **(b)** by a second lab. Representative staining for **(c-e)** nucleocapsid expression and **(f-h)** MxA expression in caudal (lower) lung of RMs necropsied at 4 dpi and 7 dpi. **(i)** Scoring for individual parameters of lung pathology of RMs necropsied at 4 dpi and 7 dpi. Untreated animals are depicted in red and IFNmod treated animals are depicted in blue. Black lines represent the median viral load or score of animals from each respective treatment group. Statistical analyses were performed using non-parametric Mann-Whitney tests. * p-value < 0.05, ** p-value < 0.01.

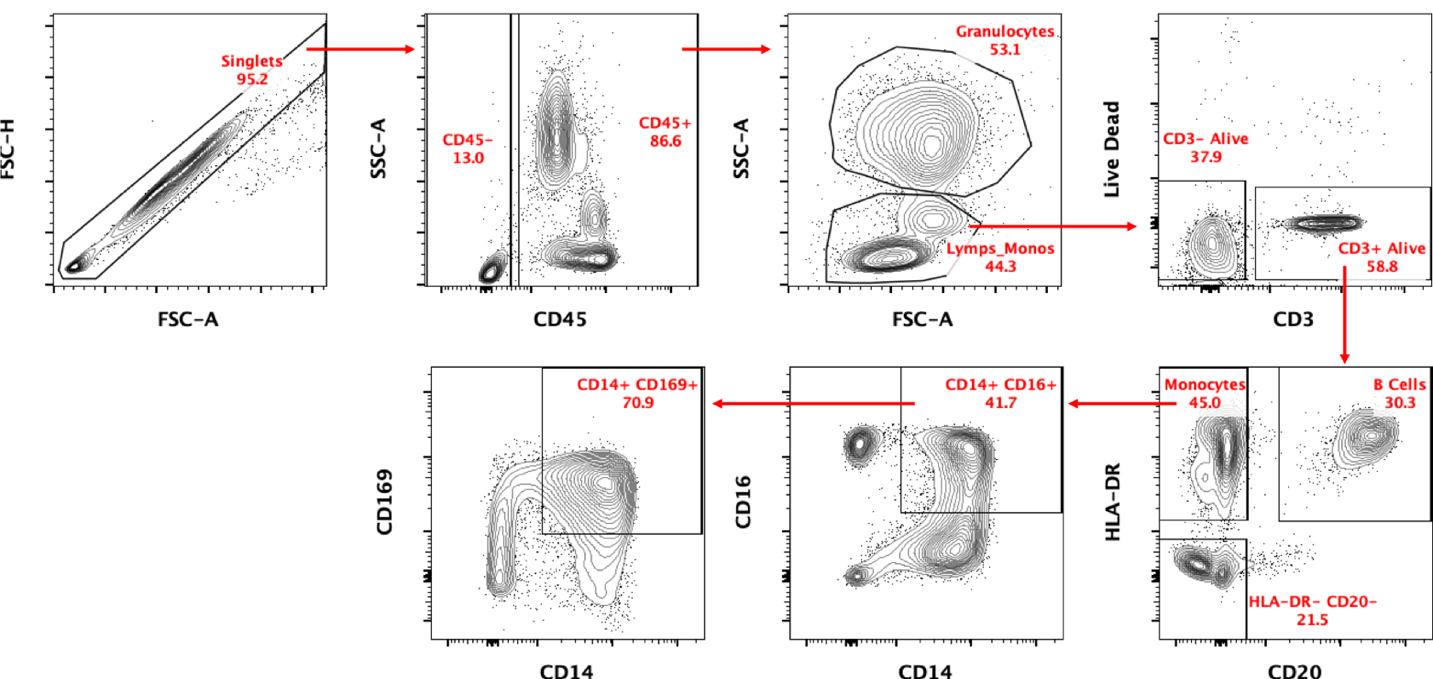
**Supplementary Fig. 5. Flow gating strategy for innate immune cell phenotyping panel used in whole blood and BAL.**

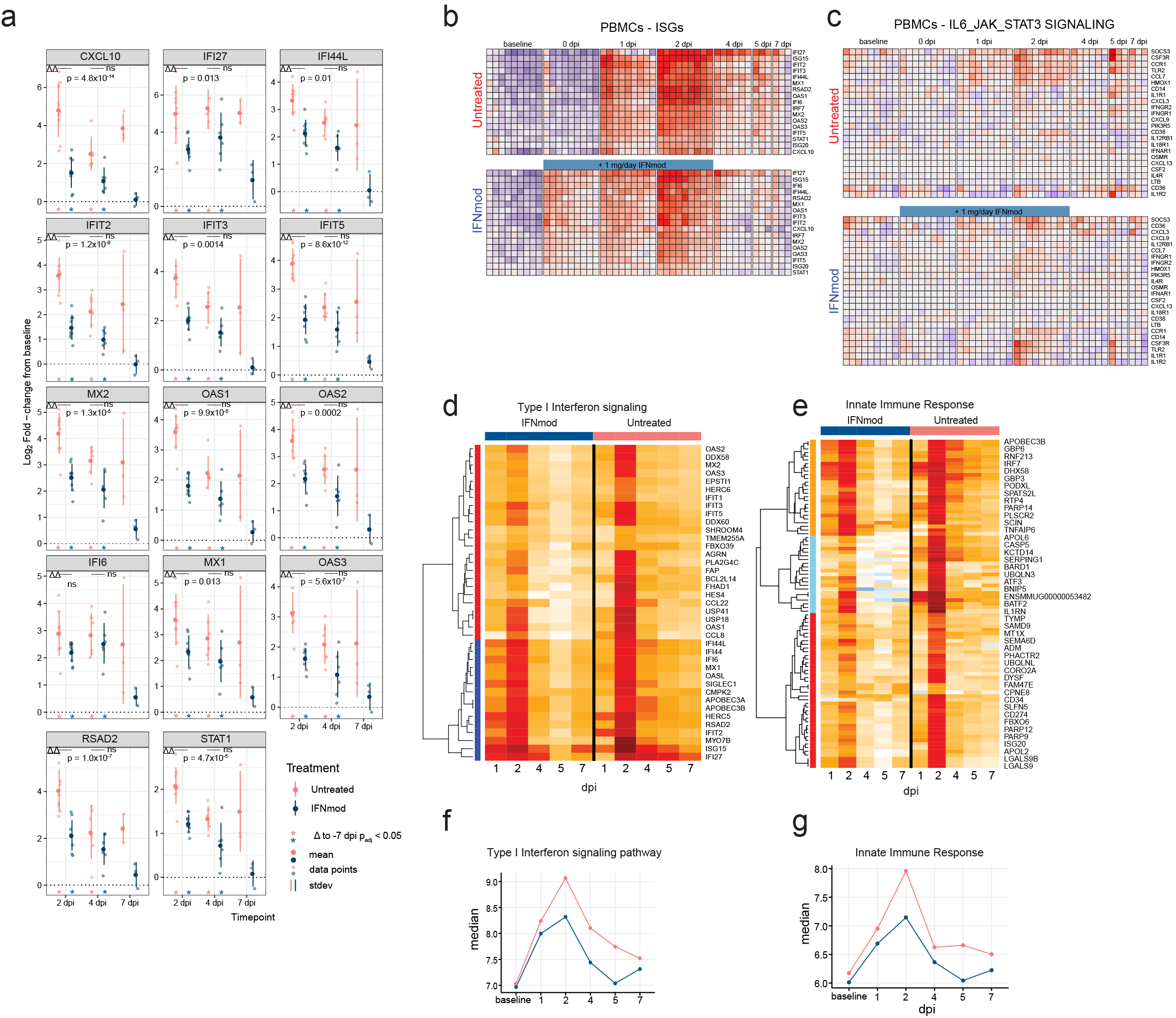

**Supplementary Fig. 6. Gene expression in BAL, PBMCs, and whole blood in rhesus macaques infected with SARS-CoV-2 after treatment with IFNmod. (a)** Log2 fold-changes in interferon stimulated genes (ISGs) in the BAL of SARS-CoV-2 infected animals receiving IFNmod treatment. Plots depict distribution of log2 fold-changes of select ISGs relative to baseline. Filled dots represent the mean, and lighter dots are individual data points. Asterisks indicate statistical significance (p_adj_ < 0.05) of gene expression relative to baseline within treatment groups; black horizontal bars indicate p-values of direct contrasts of the gene expression between groups at time-points (i.e. IFNmod vs Untreated). Vertical colored bars depict the standard deviation of each distribution. Heatmaps of longitudinal gene expression in PBMCs after SARS-CoV-2 infection for the ISG gene panel **(b)** and genes in the IL-6 JAK/STAT pathway **(c)**. The color scale indicates log2 expression relative to the mean of all samples. Samples obtained while the animals were receiving IFNmod administration are depicted by a blue bar. Heatmaps depicting log-fold changes for DEGs in significantly enriched type I interferon signaling **(d)** and innate immune response **(e)** pathways for both IFNmod and untreated animals in whole blood. Median logCPM values for genes in the pathways type I interferon signaling **(f)** and innate immune response **(g)** in whole blood.

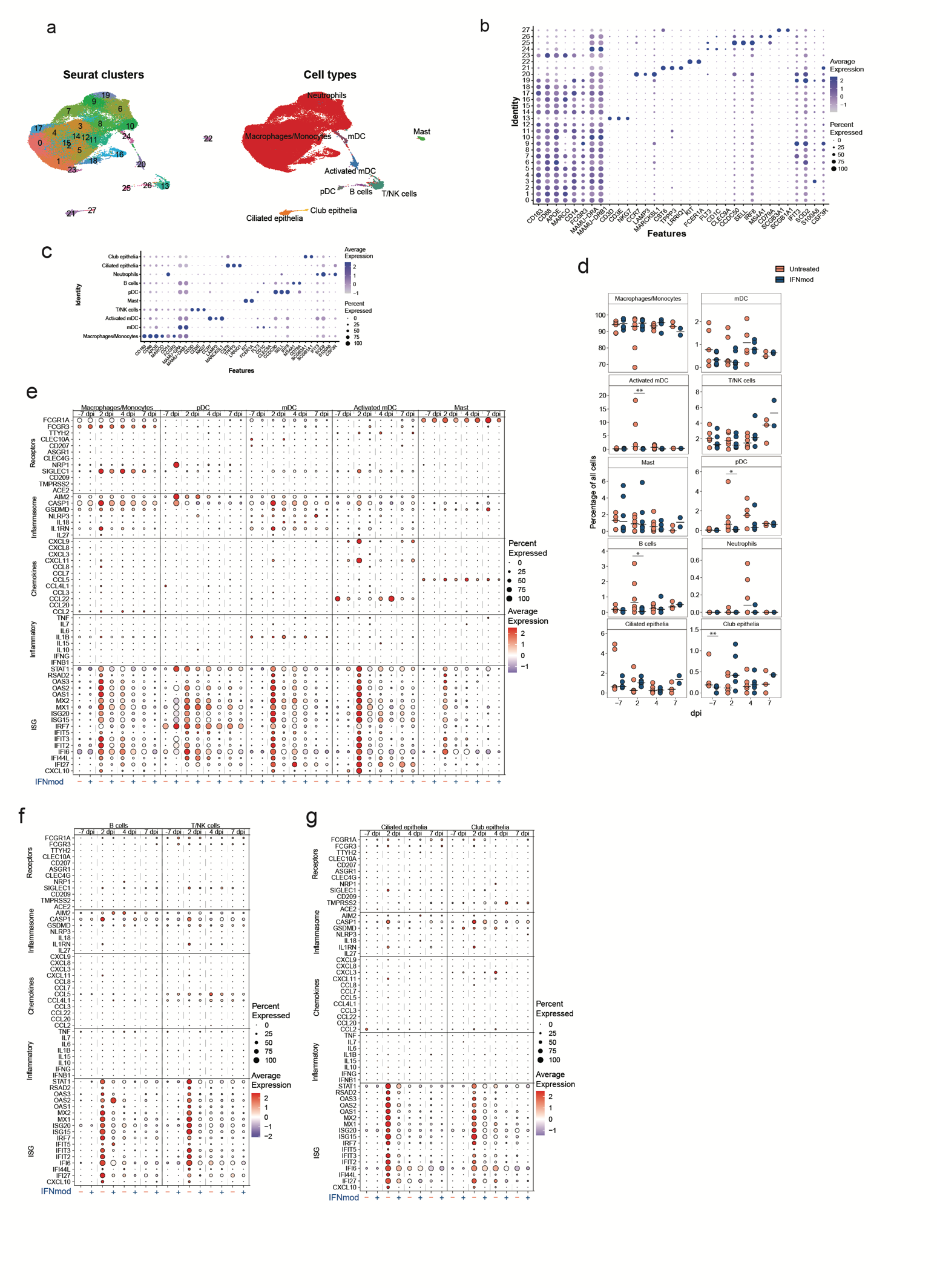

**Supplementary Fig. 7. Expression of marker genes in BAL single-cells (a)** UMAP of BAL samples colored by clusters determined using Seurat and annotated cell types. **(b)** Dot Plot showing expression of canonical marker genes in seurat clusters. **(c)** Dot Plot showing expression of canonical marker genes in annotated cell types. **(d)** Percentage of each cell type out of all BAL cells in a given sample. The black bar represents the median. Two-tailed Mann-Whitney test was used to calculate p-values. * - p-value < 0.05, ** - p-value < 0.01. **(e-g)** Dot Plot showing expression of select ISGs, inflammatory cytokines, chemokines, inflammasome-related and receptor genes in myeloid cells (neutrophils excluded due to low frequency) **(e),** lymphocytes **(f),** and epithelial cells (**g).**

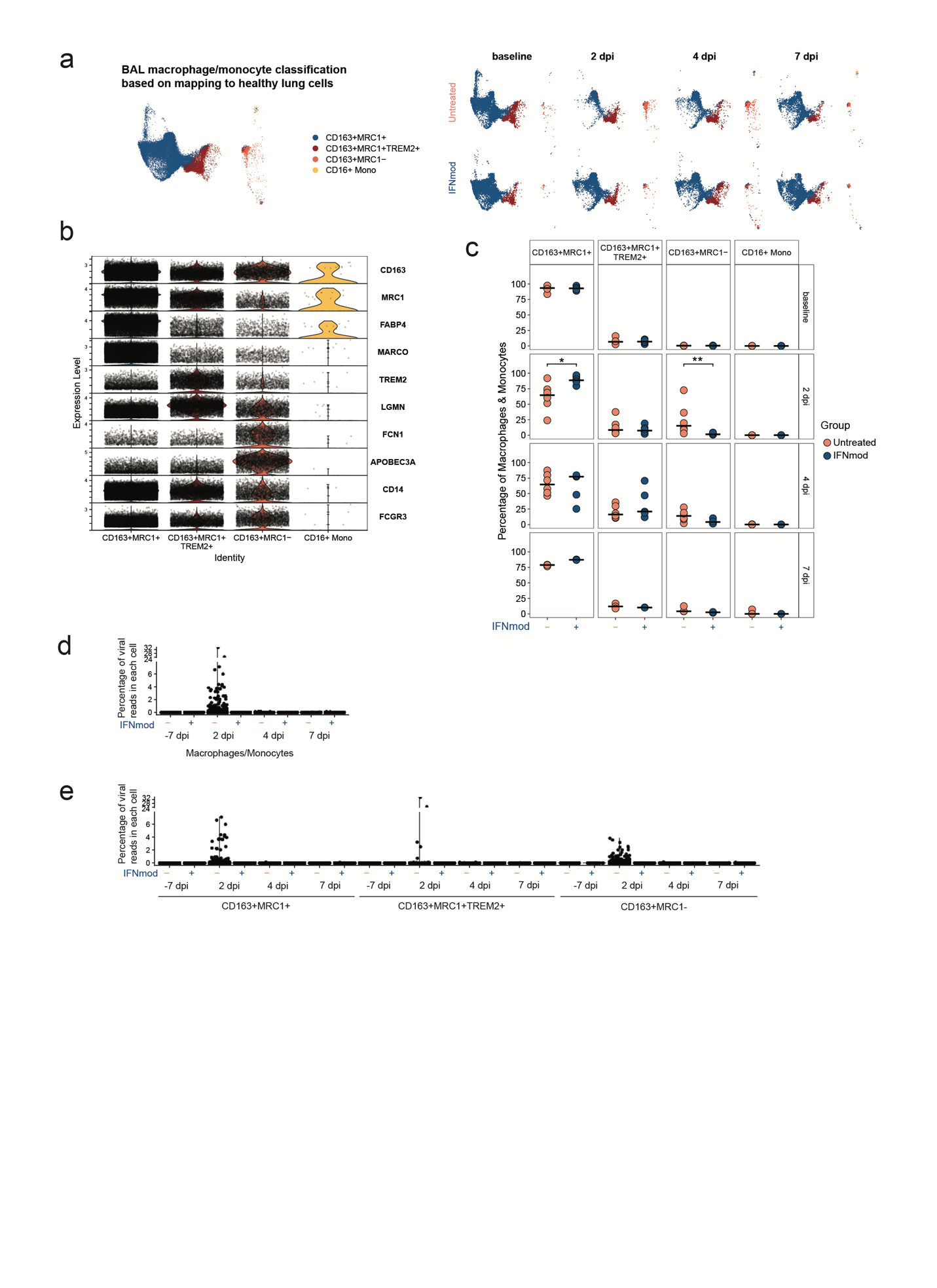

**Supplementary Fig. 8. Effect of IFNmod treatment on different BAL cell types. (a)** UMAP showing BAL macrophages/monocytes mapped to the reference macrophage/monocytes from lungs of healthy rhesus macaques. The UMAP split by time point and treatment are also shown. **(b)** Expression of marker genes for macrophage/monocyte subsets in BAL. **(c)** Percentage of different macrophage/monocyte subsets out of all the macrophages/monocytes in BAL at different time points between the untreated and IFNmod treated rhesus macaques. The black bars represent the median. Two-tailed Mann-Whitney test was used to calculate p-values. * - p-value < 0.05, ** - p-value < 0.01. Violin plots showing the percentage of viral reads in BAL total macrophages/monocytes **(d)** and individual macrophage/monocyte subsets **(e)** between the untreated and IFNmod treated macaques at different time points. The percentages were determined using the PercentageFeatureSet in Seurat for SARS-CoV2 genes. CD16+ monocytes were very excluded due to low frequency.

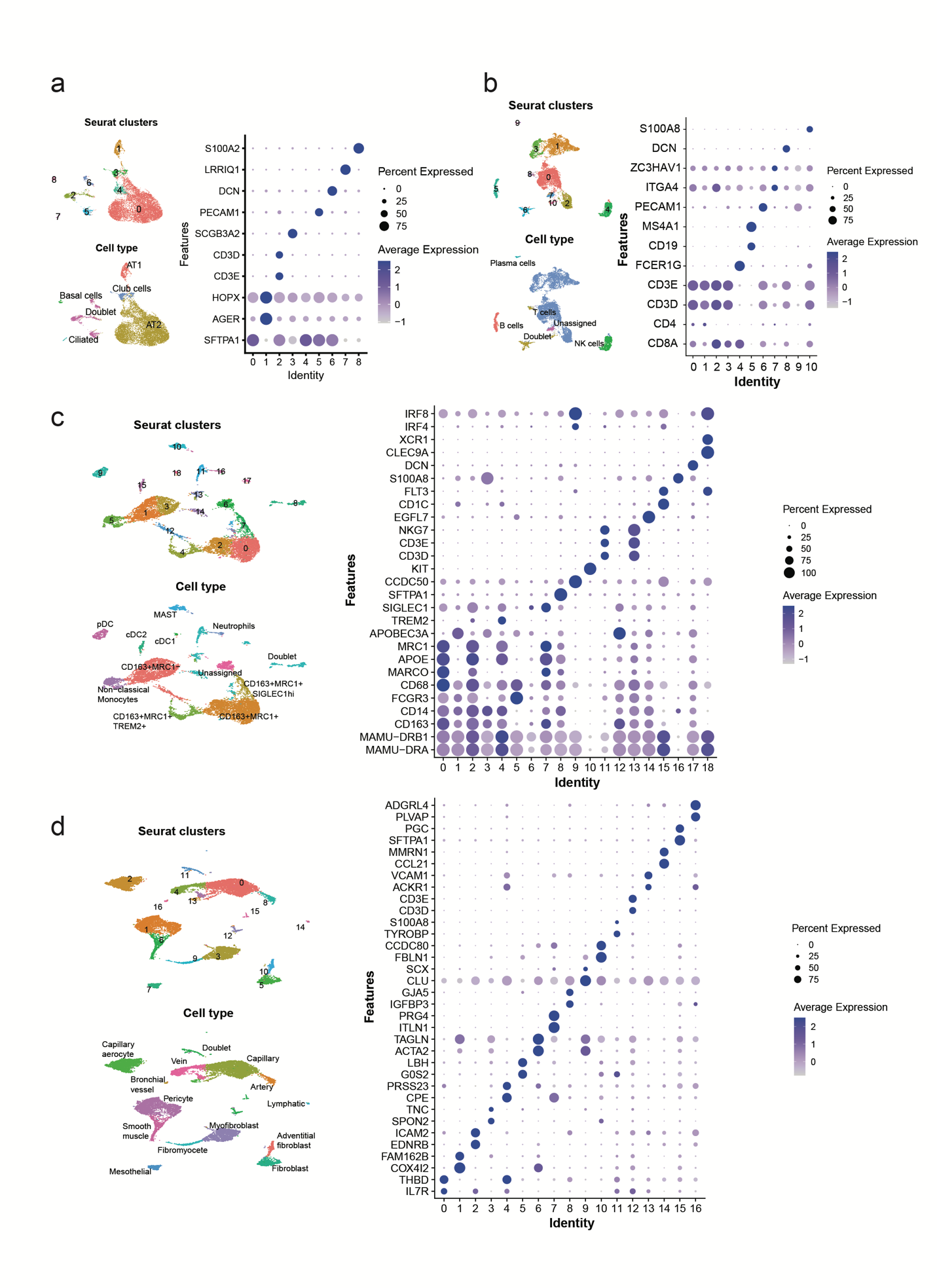

**Supplementary Fig. 9. Cell annotation of lung samples.** The cells were divided into four major categories - epithelial **(a),** lymphoid **(b),** myeloid **(c),** and other (stromal and endothelial) **(d)** and clustered separately. The clusters so obtained were annotated based on the expression of canonical markers. For each category, a UMAP and a DotPlot with canonical marker genes are shown based on seurat clustering. The second UMAP shows the cell type annotations based on the expression of marker genes.

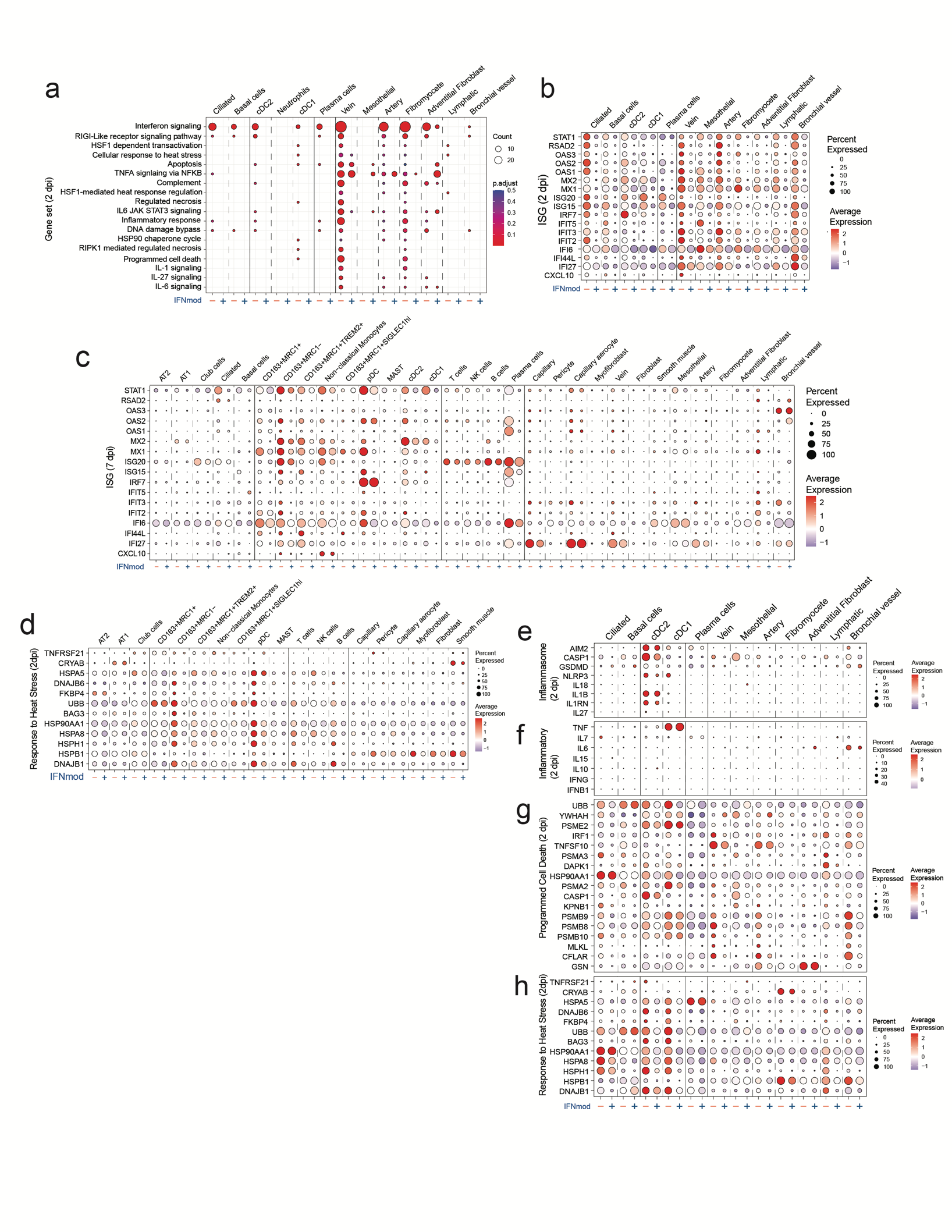

**Supplementary Fig. 10. Effect of IFNmod treatment on gene expression of lung cells.** **(a)** Enrichment of gene sets in lung cell types that were present at low frequency at 2 dpi based on over-representation analysis using Hallmark, Reactome, KEGG, and BioCarta gene sets from msigdb. Selected gene sets that were enriched in untreated RM samples at 2 dpi (p-adjusted value < 0.05) are shown. The size of the dots represents the number of genes that were enriched in the gene set and the color indicates the p-adjusted value. The gene set id in order are: M983, M15913, M27255, M27253, M5902, M5890, M5921, M27250, M41804, M5897, M5932, M27698, M27251, M29666, M27436, M27895, M27897, M1014. **(b-h)** Dot plots showing gene expression in lung cells (neutrophils not shown due to low frequency) from untreated and IFNmod treated animals. **(b)** Expression of ISG in cells present at low frequencies at 2 dpi. **(c)** Expression of ISGs in all lung cells at 7 dpi. **(d)** Expression of genes related to response to heat stress in lung cells that were present at higher frequencies at 2 dpi. **(e-g)** Expression of genes related to inflammasome **(e),** inflammation **(f),** programmed cell death **(g),** and response to heat stress **(h)** at 2 dpi in lung cell types that were present at low frequencies.

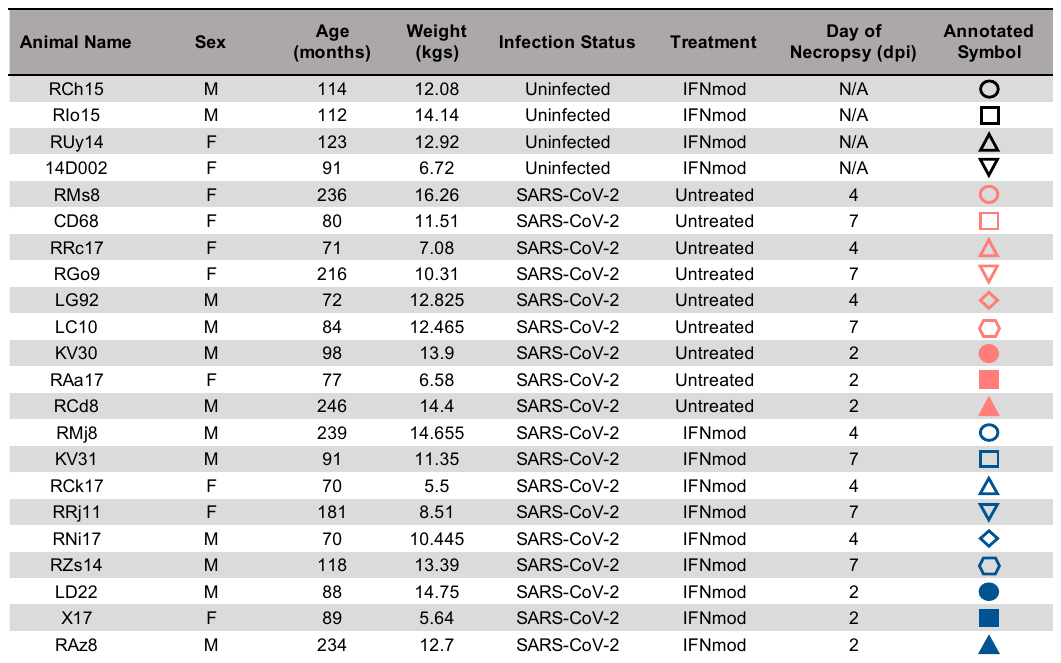

**Supplementary Table 1.** **Uninfected and SARS-CoV-2-infected macaque characteristics.** Animal ID. Sex. Age in months and weight at beginning of study in kg. Infection status. Treatment group assignment. Day post infection that necropsy was performed. Annotated symbol in figures. For infected animals, open symbols indicate that the animal was necropsied at 4 dpi or 7 dpi while filled symbols indicate that the animal was necropsied at 2 dpi.

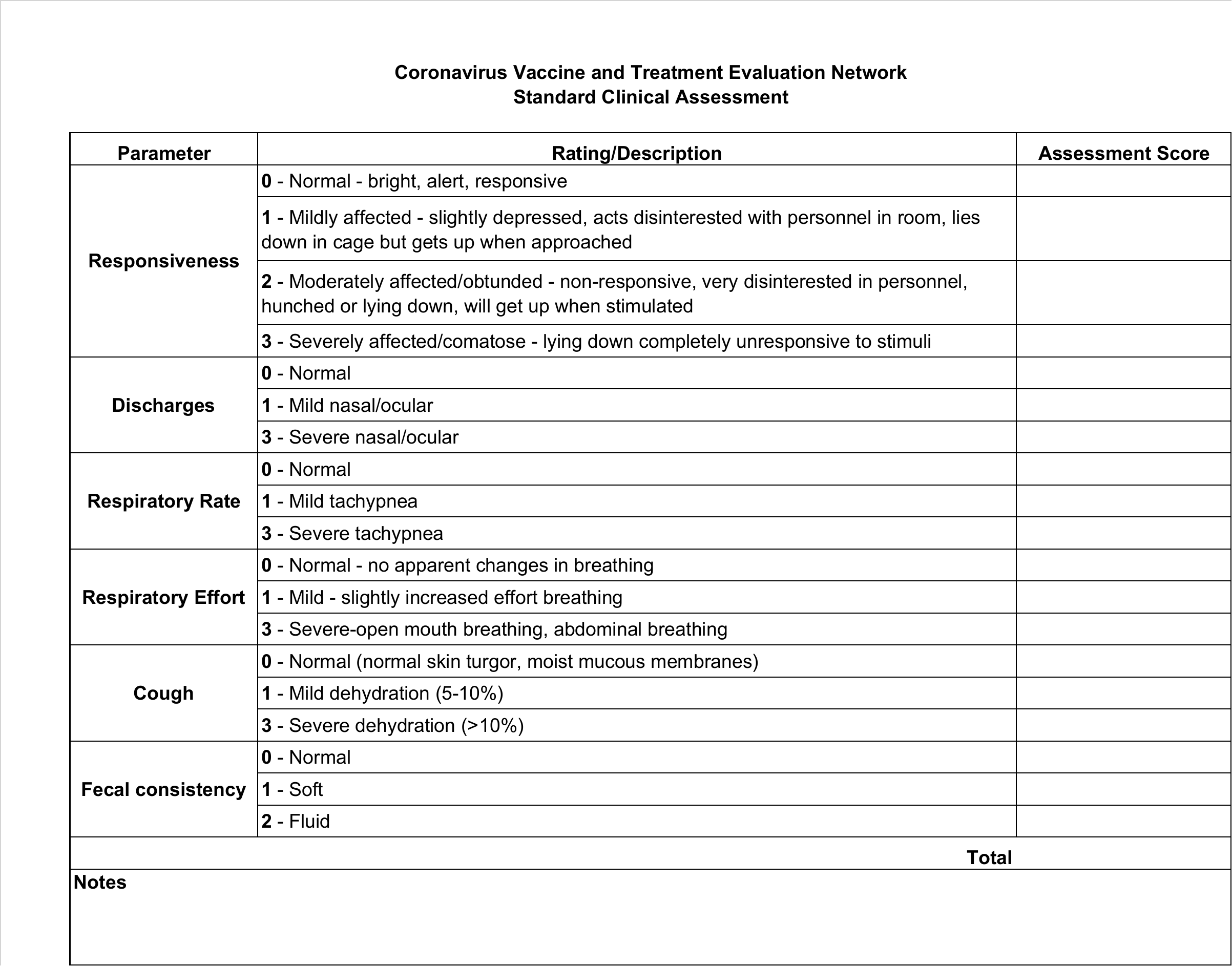

**Supplementary Table 2. Coronavirus Vaccine and Treatment Evaluation Network (CoVTEN) standard clinical assessment for cage-side scores, related to Supplementary Fig. 3b.** Cage-side scores were performed at 0, 1, 2, 3, 4, 5, and 7 dpi and added to anesthetized scores to obtain the total clinical score for each dpi. Cageside scores were based on responsiveness, discharges, respiratory rate, respiratory effect, cough, and fecal consistency and were completed prior to anesthesia.

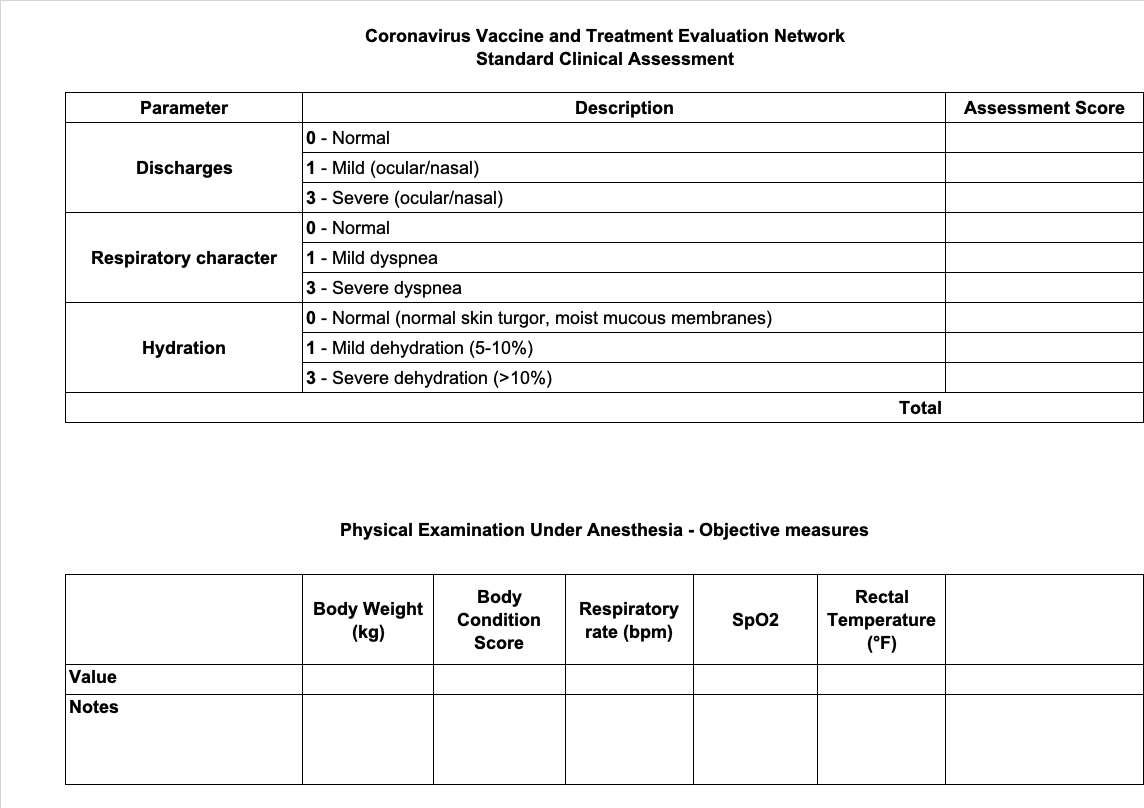

**Supplementary Table 3. Coronavirus Vaccine and Treatment Evaluation Network (CoVTEN) standard clinical assessment for anesthetized scores, related to Supplementary Fig. 3b.** Anesthetized scores were performed at 0, 1, 2, 3, 4, 5, and 7 dpi and added to cageside scores to obtain the total clinical score for each dpi. Anesthetized scores were based on discharges, respiratory character, and hydration. Body weights (kg), body condition scores, respiratory rates (bpm), SpO2 (%), and rectal temperatures (°F) were also recorded during anesthetic accesses.
